# MetaPilot enables adaptive genome-aware DDA and DIA metaproteomics using microbial genome catalogues

**DOI:** 10.64898/2026.06.12.728088

**Authors:** Kai Cheng, Daniel Figeys

## Abstract

Metaproteomics directly measures microbial protein expression in complex communities, but analysis is constrained by large, poorly structured search spaces. Microbial gene catalogues provide broad coverage, but their pooled organization hinders genome-level evidence accumulation and consistent DDA and DIA analysis. Here, we present MetaPilot, a genome-aware workflow that uses conserved marker-protein evidence to guide adaptive genome-resolved search-space refinement. MetaPilot maps identifications to candidate genomes, ranks them by marginal peptide contribution, and constructs refined sample-specific search spaces for final identification, quantification and multi-layer reporting.

Across DDA and DIA datasets from defined mixtures and faecal microbiomes, MetaPilot adapted genome selection to sample complexity, preserved substantial overlap with published peptide identifications and expanded the detectable peptide space; iterative refinement further increased DIA identification while controlling search-space expansion. In DDA-independent reanalysis of Orbitrap human gut metaproteomes, MetaPilot identified 24.4% more peptides than the published DDA-derived library and more than twice the number identified by the matched DDA-assisted workflow. In a timsTOF DIA-PASEF mouse intestinal dataset, MetaPilot outperformed uMetaP in identification depth and enabled genome-resolved functional interpretation. These results establish that DDA-independent DIA metaproteomics, guided by genome-resolved marker evidence, can exceed DDA-assisted workflows in identification depth.

## Introduction

Characterizing the functional activity of microbial communities remains a central challenge in microbiome research. Metagenomics has transformed our understanding of microbial gene content and community structure, but it primarily describes functional potential. Many microbial genes are conditionally expressed, regulated at post-transcriptional or translational levels, or contribute differently depending on community context. Metaproteomics addresses this gap by directly measuring expressed microbial proteins, providing a molecular readout of realized biological activity^1^. This makes metaproteomics particularly valuable for host-associated microbiomes, where taxonomic composition alone may not capture ecosystem function.

Human gut metaproteomics remains computationally difficult because peptide identification depends strongly on the structure and size of the protein search space^2, 3^. Comprehensive human gut gene or protein catalogues provide broad sequence coverage^4^, but they are commonly organized as pooled gene collections rather than genome-resolved units. This limits genome-level evidence accumulation and makes it difficult to interpret peptide identifications in the context of candidate organisms. Microbial genome catalogues, including metagenome-assembled genome (MAG) resources, preserve the linkage between proteins, genomes and taxonomy, providing a better structure for genome-aware search-space refinement.

A second challenge is the lack of a unified analysis solution for both data-dependent acquisition (DDA) and data-independent acquisition (DIA) metaproteomics. DDA workflows can search broad FASTA databases for protein database refinement or direct peptide identification, although this often incurs substantial computational cost^5^. DIA analysis is less tolerant of large and complex search spaces, because the size and composition of the searched FASTA or spectral library directly influence peptide competition, FDR control and identification depth. As a result, DDA and DIA datasets are commonly processed using different tools and output structures. DIA-specific workflows often rely on pre-built or reduced spectral libraries/databases to make the search space tractable^6^, but this introduces a library-sample mismatch problem: the library may not fully represent the taxonomic and protein composition of the DIA samples being analysed. This limitation can persist even with related or matched DDA-derived libraries, because DDA sampling is less comprehensive than DIA in complex metaproteomic mixtures. Together, these issues restrict peptide detection and hinder one-stop analysis that produces harmonized peptide-, protein-, genome-, taxonomic- and functional-level outputs across acquisition modes^7, 8^.

Genome-resolved catalogues therefore provide a practical foundation for adaptive metaproteomic search-space construction. Large-scale resources such as the MGnify Genomes portal offer biome-specific genome collections that preserve protein-to-genome and genome-to-taxonomy relationships^9, 10^. This genome-resolved structure allows conserved and frequently detectable marker proteins to be extracted from each genome and used as a compact first-pass search space. Peptide evidence from this marker layer can then be mapped back to candidate genomes before constructing a refined sample-specific search space.

Here, we present MetaPilot, a software platform providing genome-aware metaproteomics workflows for adaptive genome-resolved search-space refinement across DDA and DIA data. MetaPilot retrieves biome-specific genome databases from MGnify and uses marker-protein evidence to rank candidate genomes, select sample-specific genome sets and construct refined search spaces for final analysis. We first describe the workflow architecture and evaluate MetaPilot using paired DDA and DIA datasets from defined microbial mixtures and complex faecal samples. We then compare MetaPilot peptide evidence with published workflows to assess overlap and complementarity. Finally, we apply MetaPilot to DDA-independent reanalysis of an Orbitrap DIA human gut cohort and a timsTOF diaPASEF^11^ mouse intestinal dataset. In both, DIA metaproteomics guided by genome-resolved marker evidence recovers a deeper peptide and protein space than workflows built on matched DDA-derived spectral libraries. This reframes the DDA-derived library not as a prerequisite for deep DIA metaproteomics but as a constraint on it, and implies that the large body of public DIA data lacking matched DDA acquisition can be reanalysed to greater depth than previously accessible.

## Results

### MetaPilot establishes a unified genome-aware workflow for DDA and DIA metaproteomics

Metaproteomic peptide identification is shaped by the structure and size of the protein search space. An initial broad DDA search against a pooled catalogue may collect matched proteins for a refined database^12–14^, but this search is computationally expensive and does not naturally produce genome-level evidence. It is also poorly suited to DIA analysis, where identification depth is more sensitive to search-space size and composition.

MetaPilot addresses this limitation by using genome catalogues as the central reference structure. In a genome database, each protein remains linked to its source genome, allowing peptide evidence to be interpreted at the genome level. Extending earlier reduced marker-protein search strategies^15, 16^, MetaPilot extracts marker proteins from each genome and uses the resulting peptide evidence to rank and select candidate genomes (**Figure 1a**). The current workflow uses four marker categories: ribosomal proteins, replication/transcription-related proteins, elongation factors and aminoacyl-tRNA synthetases. Together, these Marker4 proteins provide a compact first-pass evidence space compatible with both DDA and DIA workflows. Using the Unified Human Gastrointestinal Genome (UHGG) v2.0.2 database of 4,744 representative species-level genomes, we quantified the number of Marker4 proteins per genome (**Figure 1b**), theoretical tryptic peptides from these proteins (**Figure 1c**), and distinct peptide sequences after removing within-marker-space redundancy (**Figure 1d**). These distributions show that Marker4 preserves genome-resolved peptide evidence across the reference catalogue while keeping the initial search space well below the full proteome.

**Figure 1.**
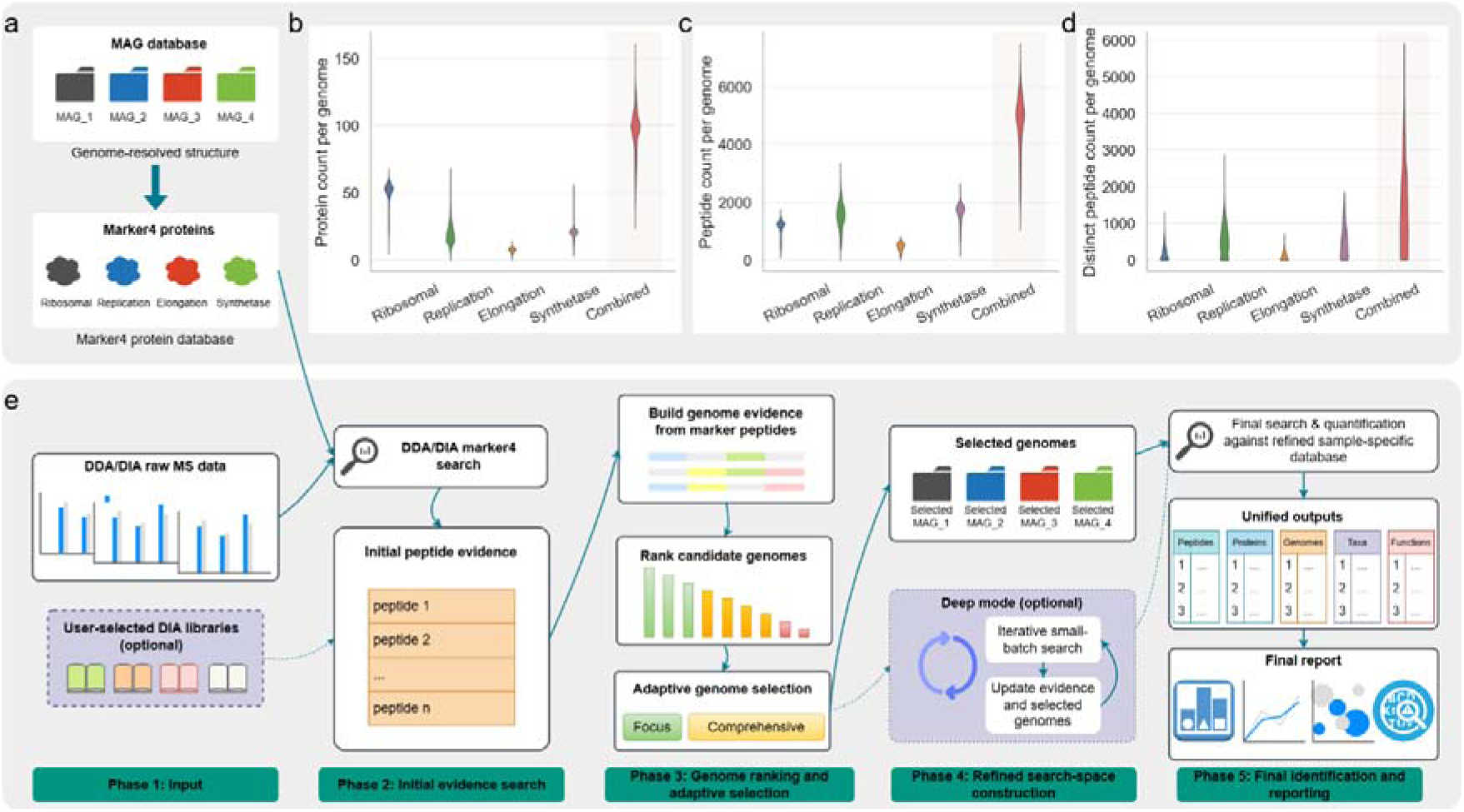
Conceptual design of the MetaPilot unified genome-aware workflow. (a) Construction of the Marker4 search space from a genome-resolved database. Proteins from four conserved and frequently detectable marker categories, ribosomal proteins, replication/transcription-related proteins, elongation factors and aminoacyl-tRNA synthetases, are extracted from each genome to generate a compact marker-protein database. **(b–d)** Distribution of marker-protein counts, theoretical peptide counts and distinct peptide counts per genome across the human gut genome database, showing that Marker4 retains genome-resolved peptide evidence while remaining substantially smaller than the full proteome search space. **e**, Overview of the unified MetaPilot workflow. DDA and DIA raw MS data, together with optional user-selected DIA libraries, are converted into initial peptide evidence using the marker-derived search space. Peptide evidence is mapped to genomes, candidate genomes are ranked, and adaptive genome selection is used to construct a refined sample-specific search space. Final DDA or DIA identification and quantification produces unified peptide-, protein-, genome-, taxonomic- and functional-level outputs with genome-level scoring and tiered reporting.

The overall workflow follows a shared genome-aware analysis logic (**Figure 1e**). DDA and DIA raw data first generate peptide evidence through their respective search engines. MetaPilot maps identified peptides to genome-derived proteins and candidate genomes, then uses genome-level evidence to rank genomes and select a sample-specific genome set. Proteins from selected genomes are used to construct the refined search space for final identification and quantification. DDA and DIA therefore share the same downstream logic for genome ranking, search-space refinement and multi-level export. For large Marker4 collections with more than 100,000 sequences, MetaPilot can further reduce the initial DIA search burden by applying DeepDetect^17^ peptide-detectability prediction and retaining high-detectability marker peptides, while preserving the same Marker4-derived genome evidence structure (Supplementary Figure S1).

To evaluate this framework, we implemented several operating modes derived from the same genome-aware backbone. DDA analyses included Marker4-based workflows and a Marker2 workflow using only ribosomal proteins and elongation factors, consistent with the marker strategy used in our previous MetaLab platform^15^. DIA analyses included Marker-based modes and Seed-library modes. These modes differ in the source of initial peptide evidence and in how broadly the refined search space is expanded, but all follow the same marker-evidence-guided genome refinement framework.

### Evaluation of MetaPilot by paired DDA and DIA datasets from both simple and complex samples

We reanalysed eight public metaproteomics datasets from two independent groups, Elo^7, 8^ and Qiao^18, 19^. These datasets included a defined 12-species microbial mixture and complex human faecal samples, each acquired by both DDA and DIA: Elo_12mix_DDA (n = 3), Elo_12mix_DIA (n = 3), Elo_faecal_DDA (n = 6), Elo_faecal_DIA (n = 6), Qiao_12mix_DDA (n = 9), Qiao_12mix_DIA (n = 9), Qiao_faecal_DDA (n = 9) and Qiao_faecal_DIA (n = 9). For each dataset, results were available from two publications: the original study reported the paired DDA and DIA analyses^7, 18^, whereas the later study reanalysed the same DIA data using an optimized DIA workflow^8, 19^. We therefore compared MetaPilot DDA results with the DDA results from the original publications, and MetaPilot DIA results with the optimized DIA results from the later publications by the same groups.

We first examined genome selection behaviour across all eight datasets (**Figure 2a, b**). The number of genomes selected for refined search was strongly dependent on sample complexity and highly consistent within each dataset type. Genome-selection trajectories for individual raw files are shown in **Supplementary Figures S2–S16**. In 12mix samples, peptide coverage curves reached a knee early, producing compact refined search spaces. Faecal samples required substantially larger genome sets before incremental coverage flattened. This pattern was reproduced across both research groups and both acquisition modes, showing that MetaPilot adapts search-space size to sample complexity rather than applying a fixed database size.

**Figure 2.**
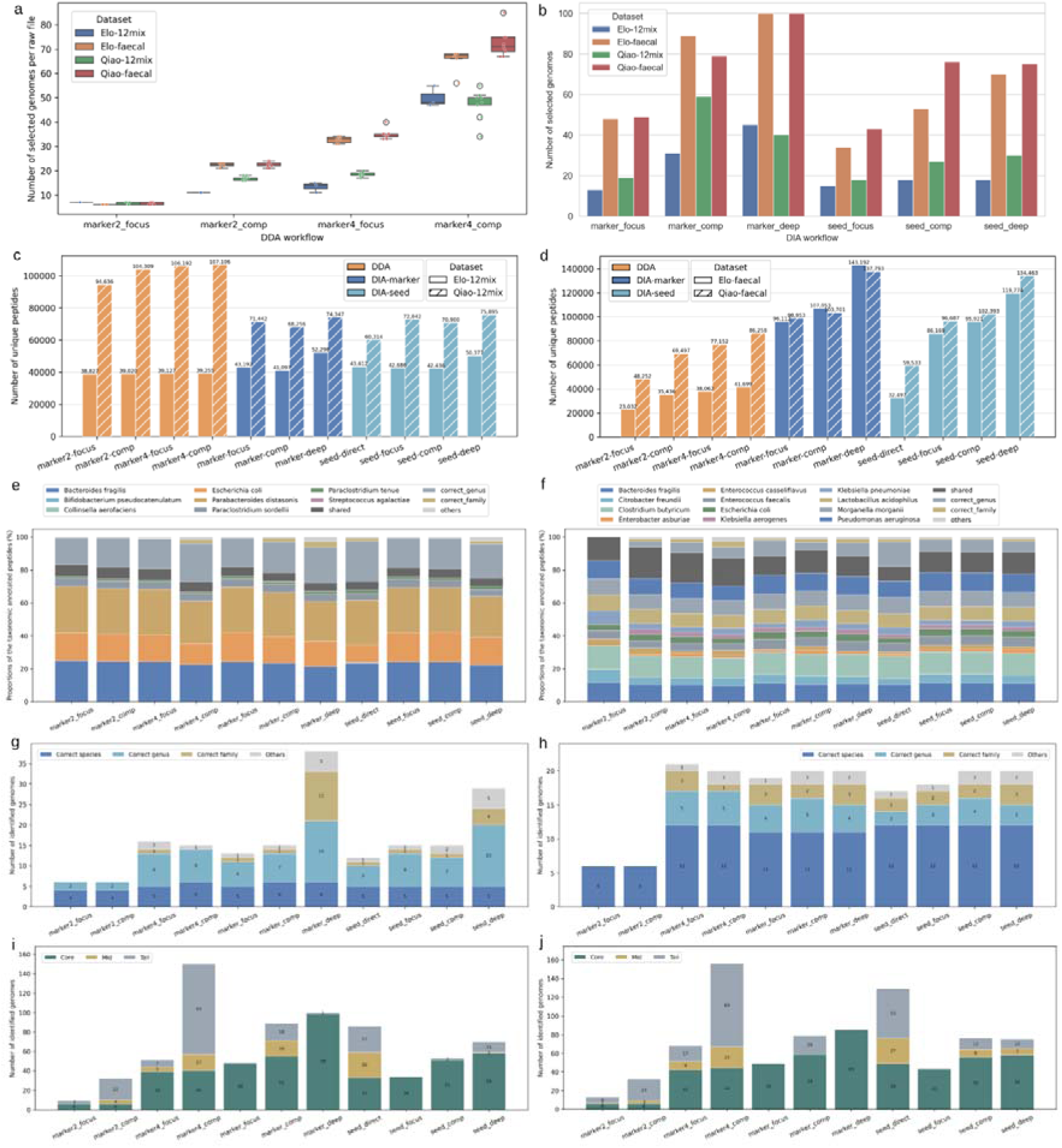
Adaptive genome refinement, peptide identification and final genome reporting across simple and complex metaproteomic benchmarks. **(a, b)** Number of genomes selected per raw file for DDA workflows **(a)** and DIA workflows **(b)** across the Elo and Qiao 12mix and faecal datasets. Focus, Comprehensive and Deep modes correspond to different stopping rules or iterative expansion strategies during genome selection. **(c, d)** Number of unique peptides identified from the two 12mix datasets **(c)** and the two faecal datasets **(d)** using representative DDA, DIA Marker and DIA Seed workflows. **(e, f)** Composition of final reported genomes in the Elo_12mix **(e)** and Qiao_12mix **(f)** datasets based on the core tier of the exported MetaPilot genome list. Genomes were classified as correct species, correct genus, correct family or others according to their agreement with the theoretical 12mix composition. **(g, h)** Composition of final reported genomes in the Elo_faecal **(g)** and Qiao_faecal **(h)** datasets, respectively, stratified into core, mid and tail tiers according to genome rank and accumulated peptide coverage.

This adaptive behaviour was reflected in peptide identification (**Figure 2c, d**). In Elo_12mix, most workflows produced similar peptide yields despite differences in the number of selected genomes, and the Deep DIA strategy provided little additional gain over Focus or Comprehensive modes (**Supplementary Data 1, 2**), indicating that peptide recovery was already close to saturation in these low-complexity samples. In Qiao_12mix, DDA produced more peptide identifications than DIA (**Supplementary Data 3**). This reflects the combination of low sample complexity, which allows DDA to approach saturated peptide recovery, and the more permissive search settings used — open search, semi-tryptic cleavage and up to two missed cleavages — which are less suited to DIA workflows where broader candidate spaces reduce sensitivity after FDR control.

In faecal datasets, DIA-based workflows clearly outperformed DDA: the deepest DDA workflow (Marker4-Comprehensive) identified 41,699 and 86,258 unique peptides in Elo_faecal and Qiao_faecal respectively, whereas the deepest DIA workflow (Marker-Deep) identified 143,192 and 137,793 (**Figure 2d, Supplementary Data 4**), consistent with the sampling advantage of DIA in high-complexity samples. In complex microbiomes, DDA top-N precursor selection becomes limiting, whereas DIA samples fragment information more comprehensively across the chromatographic space. The faecal DIA results also highlighted the importance of controlling effective search-space size. Comprehensive mode often searched nearly twice as many genomes as Focus mode but yielded only modest peptide gains (11.4% in Elo_faecal and 4.8% in Qiao_faecal), indicating that excessive DIA search-space expansion can reduce net sensitivity after FDR control. The iterative deep strategy alleviated this effect and recovered additional peptides across faecal datasets, supporting incremental genome introduction rather than single-round broad search-space expansion for complex DIA metaproteomics.

The taxonomic composition of reported genomes in the 12mix datasets supported the refinement strategy (**Figure 2e, f, Supplementary Data 1, 3**). MetaPilot exports genome lists stratified into core, mid and tail tiers based on genome ranking and accumulated peptide coverage, with the core tier representing the highest-confidence organisms. For 12mix analyses, accuracy was therefore evaluated on the core tier. Most workflows reported approximately 15 core genomes for Elo_12mix and approximately 20 for Qiao_12mix, which is reasonable when searching against the comprehensive UHGG reference rather than a custom database matched to the mixture composition. Qiao_12mix showed strong exact-species recovery, with approximately 11 of the 12 expected mixture members recovered in the core tier and few incorrect assignments.

Elo_12mix showed lower exact-species recovery than Qiao_12mix. To determine whether this reflected workflow performance or the dataset itself, we queried the original Elo study’s published peptide identifications against UniProt using Unipept^20, 21^ (**Supplementary Table 1**). Species-level evidence was confirmed for only 7 of the 12 nominal mixture members. The remaining 5 species, *Enterorhabdus sp.*, *Eubacterium tenue*, *Bifidobacterium pseudocatenulatum*, *Streptococcus agalactiae* and *Bifidobacterium bifidum*, yielded either no detectable peptide evidence or only genus-level assignments. This indicates that these species were likely absent or below the detection threshold in the actual samples. Species-level recovery is therefore not the most informative evaluation axis for Elo_12mix. For this dataset, the primary benchmark is peptide identification depth, where MetaPilot outperformed the published analysis.

In faecal datasets, where true species composition is unknown, the core, mid and tail tiers were used to inspect the final reported genome sets (**Figure 2g, h, Supplementary Data 2, 4**). Compared with the defined mixtures, faecal samples contained broader genome profiles with greater contributions from mid and tail tiers, consistent with higher community complexity and larger refined search spaces. To further assess refinement specificity under these conditions, we augmented the 4,744-genome human-gut catalogue with 562 genomes from the MGnify tomato-rhizosphere catalogue after excluding 17 genomes sharing species-level taxonomy with the target catalogue. Reanalysis of the Qiao faecal datasets retained only 1 of 562 entrapment genomes in the DDA refined search space and 2 of 562 in DIA; none reached the DDA core tier and only one reached the DIA core tier. The additional DIA survivor, GTDB-designated *Bacillus_A cereus*, remained in the tail tier and is also a species known to occur transiently in the human intestinal tract^22, 23^, whereas the sole core-tier survivor, *Pseudomonas paraeruginosa*, was a close relative of the strongly supported target-catalogue *P. aeruginosa* representative^24^. *P. paraeruginosa* was supported by 23 genome-unique peptides in DDA and 147 in DIA, none of which mapped to the ten *Pseudomonas* genomes in the target catalogue, suggesting that its retention reflected incompletely represented *Pseudomonas* sequence diversity rather than broad propagation of unrelated rhizosphere genomes (**Supplementary Figure S17, Supplementary Data 4**).

We next compared representative MetaPilot workflows with published methods using peptide-level overlap analysis (**Figure 3a–h**). One representative MetaPilot mode was selected per dataset (**Methods**). Across all datasets, MetaPilot reproduced the main peptide evidence reported by established methods while contributing complementary identifications, particularly in faecal datasets.

**Figure 3.**
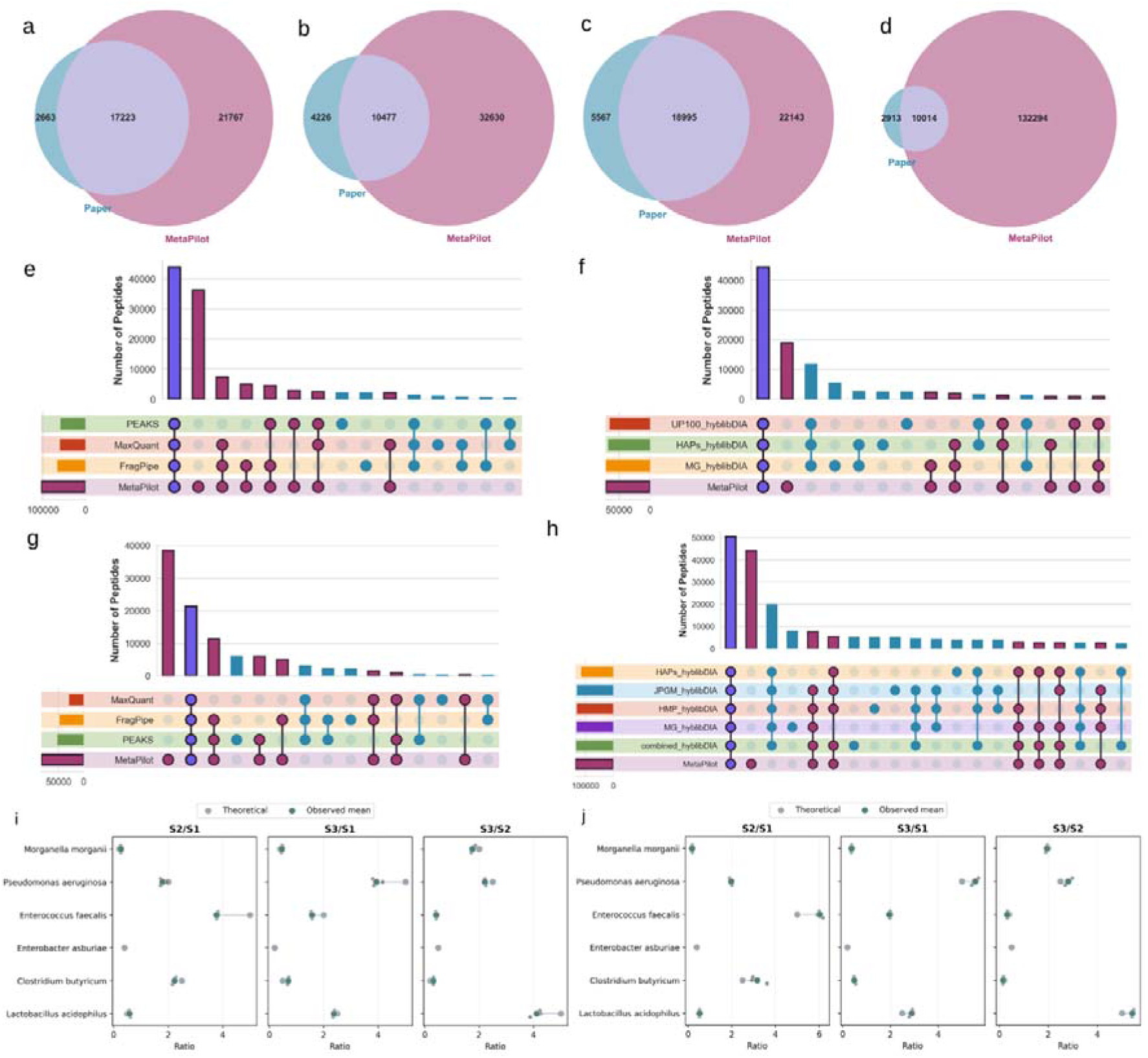
DDA and DIA benchmarks compared with published workflows. **(a–d)** Venn diagrams showing peptide overlap between MetaPilot and the corresponding published workflows for Elo_12mix_DDA, Elo_12mix_DIA, Elo_faecal_DDA and Elo_faecal_DIA, respectively. (**e–h)** UpSet plots showing peptide overlap between MetaPilot and published workflows for Qiao_12mix_DDA, Qiao_12mix_DIA, Qiao_faecal_DDA and Qiao_faecal_DIA, respectively. For these comparisons, one MetaPilot workflow was chosen per dataset to represent the most appropriate operating mode: Marker4-Focus for simple DDA, Marker4-Comprehensive for faecal DDA, Marker-Focus for simple DIA and Marker-Deep for faecal DIA. The compared published workflows differ in their underlying search spaces, including conventional DDA search engines, metagenomic-sequencing-derived databases, HAP-guided refined databases, public microbiome protein databases and hybrid DDA/DIA spectral-library strategies. **(i, j)** Quantitative evaluation of MetaPilot on the Qiao spike-in faecal benchmark, showing theoretical and observed abundance ratios for the six spiked bacterial species across pairwise sample comparisons. Observed values are shown as mean estimates across replicates.

The published workflows used different database and library strategies, which is important for interpreting the overlap results. The Elo DIA workflows used DDA-derived spectral libraries and later DIA-only pseudospectral libraries. The Qiao studies evaluated directDIA^25^, DDA-library Spectronaut^26^, DIA-NN^27^ predicted-library workflows and HAP-guided (high-abundance protein) hybrid-library strategies, termed hyblibDIA (**Supplementary Table 2**). In hyblibDIA, DIA data were analysed using libraries supported by fractionated DDA evidence after HAP-guided database refinement. MetaPilot Marker-Deep achieved peptide identification depth comparable to the combined_hyblibDIA result without requiring pooled DDA data for hybrid-library construction (**Figure 3h**).

The Qiao spike-in faecal benchmark further supported the quantitative validity of MetaPilot refinement. Proteins from six cultured species were spiked at defined relative abundances into a faecal background. MetaPilot-derived abundance ratios were consistent with the expected spike-in ratios in direction across all pairwise comparisons, and within two-fold of the expected magnitude in the majority of cases (**Figure 3i, j**), indicating that adaptive search-space refinement preserved quantitative fidelity.

### DDA-independent reanalysis of Orbitrap DIA human gut metaproteomes

The benchmark analyses above showed that DIA recovered more peptides than DDA in complex faecal microbiomes. This suggests that DDA-derived spectral libraries, which reflect the peptide space sampled by DDA, may not provide the optimal search foundation for DIA metaproteomics. To test whether a fully DDA-independent strategy could recover deeper and biologically coherent profiles, we reanalysed the Orbitrap Exploris 480 DIA data from the metaExpertPro study, comprising 78 raw files^28^. In the original workflow, metaExpertPro constructs a spectral library from DDA data and uses it for DIA-based peptide and protein quantification. MetaPilot was applied here in DDA-independent Marker-Deep mode. This mode was selected to maximize peptide recovery in the complex faecal DIA cohort by iteratively expanding the refined search space through ranked genome batches prior to the final quantifying search.

During iterative refinement, MetaPilot progressively accumulated peptide evidence, reaching 400,116 cumulative peptides by round 39 (195 genomes), where the plateau stopping criterion was satisfied (**Figure 4a**). The number of newly identified peptides decreased across successive rounds, indicating that the iterative search was approaching saturation and that further genome addition would yield diminishing returns. This evidence-guided stopping behaviour ensures that search-space expansion is driven by observed peptide gain rather than a fixed genome count, preventing unnecessary enlargement of the final library.

**Figure 4.**
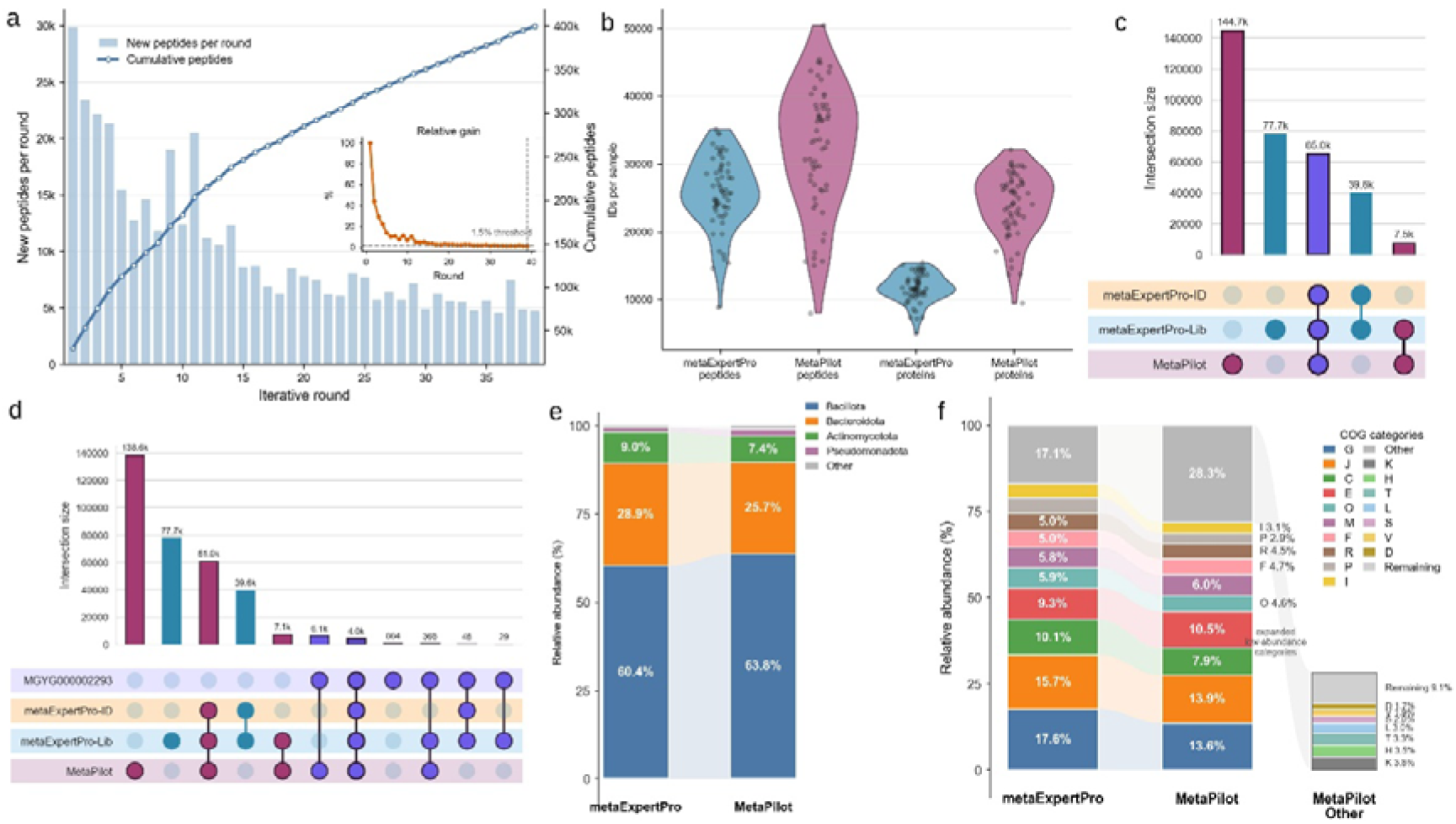
DDA-independent reanalysis of Orbitrap DIA human gut metaproteomes using MetaPilot. **(a)** Iterative MetaPilot refinement showing newly identified peptides per round and cumulative peptide identifications. The iterative search reached a plateau at round 39, where the stopping criterion was satisfied. **(b)** Comparison of peptide and protein identification depth between metaExpertPro and MetaPilot. **(c)** Peptide-level overlap among the metaExpertPro DDA-derived spectral library, metaExpertPro DIA identifications and MetaPilot DIA-independent identifications. **(d)** Constrained single-genome validation using MGYG000002293 (*Prevotella sp900557255* / *Segatella copri*) as the search space, showing peptide overlap among the constrained search, the metaExpertPro DDA-derived library, metaExpertPro DIA identifications and MetaPilot DIA-independent identifications. **(e)** Phylum-level taxonomic composition in the published metaExpertPro analysis and MetaPilot reanalysis. **(f)** COG-category composition in the published analysis and MetaPilot reanalysis, with decomposition of the MetaPilot “Other” fraction into lower-abundance functional categories.

The accumulated peptide evidence was used for the final quantifying search, from which MetaPilot identified 231,869 microbial peptides and 69,393 proteins (**Supplementary Data 5, 6**). A separate host database search yielded an additional 4,318 peptides and 857 proteins, giving totals of 236,187 peptides and 70,250 proteins (**Supplementary Data 7**). For comparison, the original metaExpertPro analysis, performed on Orbitrap Exploris 480 data, reported a DDA-derived spectral library containing 189,808 peptides and 51,269 protein groups. Applying this library to the DIA data yielded 114,571 peptides and 36,548 proteins. Both metaExpertPro values are likewise reported as totals including host sequences, so all comparisons below are made on a total-peptide basis. On this basis MetaPilot identified 24.4% more peptides than the DDA-derived library and 2.06-fold more than those recovered by the DDA-assisted DIA search. The protein-level gain was of similar magnitude: 70,250 proteins against 51,269 protein groups in the DDA-derived library (37.0% more), and against 36,548 proteins recovered by the DDA-assisted DIA search (1.92-fold more). At the per-sample level, metaExpertPro quantified on average 22,460 ± 4,964 microbial peptides and 11,301 ± 2,172 microbial protein groups, whereas MetaPilot quantified 32,377 ± 1,181 peptides and 24,023 ± 618 proteins from the same cohort. MetaPilot also exported standardized quantitative tables across all analytical layers, comprising 195 genome-resolved entries, 68,495 annotated protein rows, 51,969 NOG-level function entries and 3,305 KO-level function terms. For each sample, 1,560 ± 214 COGs and 1,640 ± 265 KOs were obtained from MetaPilot (**Supplementary Data 8**), representing increases of 13.5% and 16.6%, respectively, compared with the metaExpertPro result (1,374 ± 246 COGs and 1,406 ±109 KOs). The final total is lower than the 400,116 peptides accumulated across the 39 iterative rounds because the reported identifications derive from a single search against the final refined search space, under one round of FDR control, rather than from the union of evidence collected across rounds. All numbers reported here are therefore based on the more conservative single-search estimate.

We next compared the peptide evidence spaces of the two approaches (**Figure 4c**). The metaExpertPro DDA-derived library, the metaExpertPro DIA identifications and the MetaPilot identifications showed substantial overlap, demonstrating that MetaPilot recovered a large fraction of the peptide space supported by the DDA-assisted workflow. MetaPilot additionally identified a set of peptides absent from the metaExpertPro results, indicating that the DDA-independent workflow extended rather than simply replicated the published evidence space.

To assess the plausibility of MetaPilot-specific identifications, we performed a constrained single-genome validation. MGYG000002293 was selected because it was among the highest-confidence core-tier genomes reported by MetaPilot in this cohort. All DIA data were searched against only MGYG000002293, annotated as *Prevotella sp900557255* in MGnify and corresponding to *Segatella copri* under current taxonomy, allowing peptide evidence for this organism to be evaluated in a simplified search space (**Figure 4d, Supplementary Data 9**). Approximately 4,002 peptides were shared across all four comparison sets, confirming a robust species-level signal. MetaPilot additionally identified 6,112 peptides not detected by metaExpertPro. To assess taxonomic specificity, peptides were queried against UniProt using Unipept. Among the shared peptides, 25.6% (1,024/4,002) were specifically assigned to *Segatella copri* (**Supplementary Table 3**). Among MetaPilot-specific peptides, this fraction increased to 40.7% (2,489/6,112, **Supplementary Table 4**). The enrichment of species-specific assignments among MetaPilot-exclusive peptides supports their origin from a well-represented organism. Furthermore, although the standard MetaPilot analysis searched a comprehensive microbiome database rather than this single-genome search space, it recovered 90.1% of the peptides observed in the constrained search, indicating that broader database searching did not substantially compromise sensitivity for a strongly supported genome.

We then examined whether the increased identification depth preserved broad biological structure. At the phylum level, MetaPilot reproduced the dominant taxonomic composition of the original analysis, with *Bacillota* and *Bacteroidota* as the two major phyla (**Figure 4e**). *Bacillota* accounted for 63.8% in the MetaPilot profile versus 60.4% in the published result; *Bacteroidota* accounted for 25.7% versus 28.9%. *Actinomycetota* and *Pseudomonadota* were recovered at comparable levels. The additional identifications obtained by MetaPilot therefore did not distort the broad taxonomic structure of the dataset.

A consistent pattern was observed at the functional level (**Figure 4f**). The major COG categories in the MetaPilot profile — carbohydrate transport and metabolism, translation, energy production and conversion, and amino acid transport and metabolism — were broadly concordant with the published result. MetaPilot produced a larger aggregated “Other” fraction, which on decomposition comprised multiple lower-abundance but interpretable categories, including transcription, coenzyme transport and metabolism, signal transduction, replication and repair, defence mechanisms and cell cycle-related functions. The expanded “Other” fraction therefore reflects increased coverage of lower-abundance functional categories rather than disagreement in the dominant functional structure.

Together, these results demonstrate that MetaPilot recovers deep and biologically coherent metaproteomic profiles from Orbitrap DIA data without DDA-derived spectral libraries. These findings support MetaPilot as a practical DDA-independent strategy for genome-aware reanalysis of public and cohort-scale DIA metaproteomic datasets.

### Comparison on a published timsTOF DIA-PASEF mouse intestinal dataset

uMetaP was introduced as a DIA-PASEF metaproteomics workflow for microbial peptide and protein quantification^29^. Its database strategy uses novoMP, which expands the searchable metaproteomic database with *de novo* peptide sequences obtained from DDA-PASEF spectra followed by homology-based protein retrieval. Thus, uMetaP is best viewed as a DIA-centred workflow supported by DDA-derived *de novo* evidence. To evaluate MetaPilot against this framework, we reanalysed the colonic-content dataset used in the original uMetaP study. The dataset comprises samples from control Hsp60^fl/fl^ mice and intestinal epithelial cell-specific Hsp60^Δ/ΔIEC^ mice collected at day 0 (D0) and day 8 (D8), yielding four groups: fl/fl_D0 and fl/fl_D8 as controls, and Δ/ΔIEC_D0 and Δ/ΔIEC_D8 as early and late stages of the epithelial injury model.

MetaPilot identified more peptides than uMetaP in all four groups (**Figure 5a, Supplementary Data 10**): 125,545 versus 73,076 in fl/fl_D0, 151,224 versus 83,582 in fl/fl_D8, 77,335 versus 54,522 in Δ/ΔIEC_D0, and 87,857 versus 39,982 in Δ/ΔIEC_D8, corresponding to increases of 71.8%, 80.9%, 41.8% and 119.7%, respectively. This difference likely reflects the advantage of MetaPilot’s sample-adaptive search-space refinement, which uses peptide evidence from the analysed DIA dataset itself, whereas uMetaP relies on a pre-built *de novo*-expanded mouse faecal database generated from separate fractionated DDA-PASEF data and therefore may not fully match the composition of the Hsp60 colonic-content samples. The complete MetaPilot reanalysis produced 341,216 peptides, 76,435 proteins, 120 genomes, 76 genera, 3,361 KEGG KOs, 1,413 EC terms, 2,791 COG accessions, 59,485 NOG accessions, 4,531 GO terms, 61,837 host peptides and 5,956 host proteins (**Figure 5b, Supplementary Data 10–13**). This output depth enabled the dataset to be examined across microbial, functional and host-associated layers rather than by peptide counts alone.

**Figure 5.**
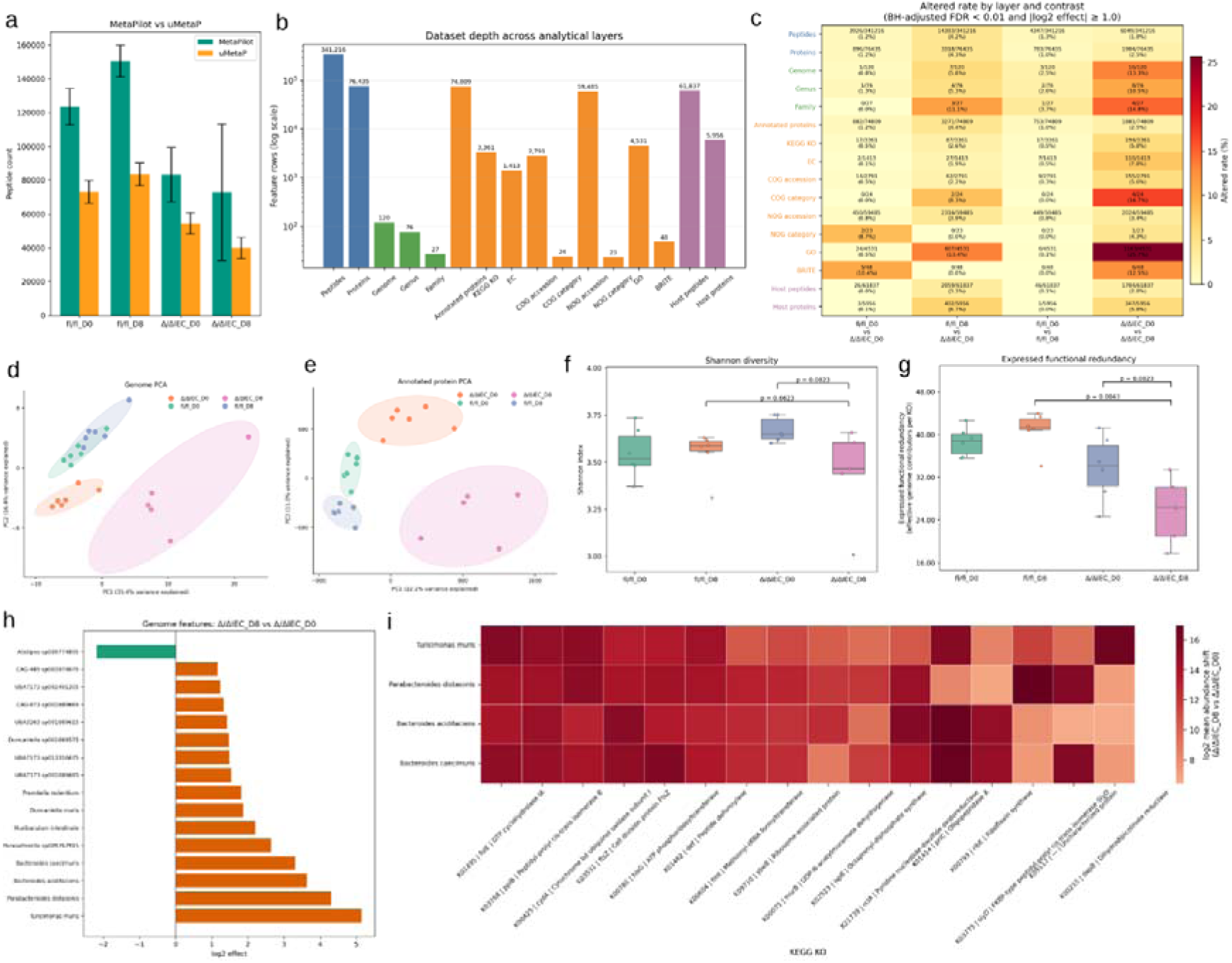
End-to-end comparison of MetaPilot on a published mouse intestinal DIA metaproteomics dataset. **(a)** Peptide identifications obtained by MetaPilot and uMetaP across the four colonic-content groups. **(b)** MetaPilot output depth across microbial peptide, microbial protein, genome, taxonomic, microbial functional and host protein layers. **(c)** Altered feature rates across analytical layers and pairwise comparisons. For each layer, features were compared between predefined groups using a two-sided Mann– Whitney U test, followed by Benjamini–Hochberg correction within that layer. Altered features were defined as BH-adjusted FDR < 0.01 and |log2 fold change| ≥ 1. **(d, e)** PCA of genome-level profiles **(d)** and annotated-protein profiles **(e)**. **(f, g)** Shannon diversity **(f)** and expressed functional redundancy **(g)** across the four groups. Displayed pairwise comparisons were performed using a two-sided Mann–Whitney U test. Functional redundancy was calculated as the effective number of genome contributors per KEGG KO. **(h)** Selected altered genomes in the Δ/ΔIEC_D8 versus Δ/ΔIEC_D0 comparison. **(i)** Shared KEGG KO trends across four concordantly increased core genomes.

The multi-layer altered-feature profile showed that the strongest perturbation was associated with the injured Δ/ΔIEC_D8 state (**Figure 5c**). Injury-associated comparisons yielded more altered features than the control timepoint comparison at both taxonomic and expressed-function levels, with the clearest signal observed at the expressed-function and annotated-protein levels. Genome-level PCA captured condition-associated structure, whereas annotated-protein PCA separated the four groups more clearly (**Figure 5d, e**). This indicates that the expressed metaproteome provided a sharper readout of biological state than genome abundance alone, and that the increased identification depth translated into improved functional resolution.

Ecological summaries supported this interpretation. In Δ/ΔIEC_D8, peptide and protein identifications were lower than in the corresponding control group, and Shannon diversity showed only a modest decrease (**Figure 5f**). Shannon diversity showed a non-significant decrease from Δ/ΔIEC_D0 to Δ/ΔIEC_D8 (p = 0.0823, Mann–Whitney U test), while the Δ/ΔIEC_D8 versus fl/fl_D8 comparison did not reach significance (p = 0.6623). By contrast, expressed functional redundancy decreased more clearly (**Figure 5g**). Functional redundancy was significantly lower in Δ/ΔIEC_D8 than in fl/fl_D8 (p = 0.0043). It was also lower in Δ/ΔIEC_D8 than in Δ/ΔIEC_D0 at the mean level, although this comparison was not statistically significant (p = 0.0823). This indicates that the same KEGG functions were supported by fewer effective genome contributors in the injured D8 state. Thus, the injured D8 samples were characterized not simply by reduced alpha diversity, but by a stronger loss of expressed functional buffering at the metaproteome level.

The genome-level differential result added an important layer to this pattern (**Figure 5h**). Despite lower peptide depth, lower alpha diversity and lower expressed functional redundancy in Δ/ΔIEC_D8, the most significantly altered genomes were predominantly increased rather than decreased. Several genomes including *Turicimonas muris*, *Parabacteroides distasonis*, *Bacteroides acidifaciens* and *Bacteroides caecimuris* showed increased expression during injury progression, whereas *Alistipes sp009774895* decreased. The injured D8 metaproteome was therefore not simply losing community complexity; rather, a subset of core genomes became more prominent within a less diverse and less redundant expressed ecosystem.

To connect these genome-level shifts to expressed function, we examined shared KEGG KO trends across the four concordantly increased core genomes (**Figure 5i**). These genomes shared functional signatures spanning cofactor metabolism, redox and respiratory function, cell division, cell envelope biosynthesis, translation and protein processing. This genome-to-function linkage completes an interpretive chain across the full analytical depth of MetaPilot: the same condition that showed reduced peptide depth, reduced alpha diversity and reduced expressed functional redundancy also exhibited selective expansion of specific genomes with coherent shared functional profiles. Together, this analysis shows that MetaPilot improves DDA-independent DIA metaproteomics at three connected levels: identification depth, expressed-function stratification and genome-resolved functional interpretation.

## Discussion

This study presents MetaPilot as a genome-aware strategy for controlling metaproteomic search spaces across DDA and DIA analyses. The central contribution is the systematic conversion of initial marker-peptide evidence into a sample-specific genome search space. This design allows MetaPilot to use comprehensive genome catalogues without requiring each analysis to search the full protein space.

The genome-resolved structure of the database is essential to this workflow. MetaPilot retains the link between each protein and its source genome, allowing Marker4 evidence to be accumulated at genome level. Candidate genomes can then be ranked by peptide support, selected adaptively and used to construct refined sample-specific databases. The four Marker4 categories provide a compact entry point into the reference catalogue while retaining genome-level information. The key contribution is therefore not simply the use of marker proteins, but their integration into a genome-ranking, adaptive-selection and refined search-space construction strategy.

The paired Elo and Qiao benchmarks showed that MetaPilot adapts its search space to sample complexity. Defined mixtures required compact genome sets and reached near-saturated peptide recovery, whereas complex faecal samples required broader genome sets and showed a stronger advantage for DIA. These results support a central design principle of MetaPilot: the refined search space should be determined by sample-specific peptide evidence rather than by a fixed database size. The Qiao spike-in benchmark showed that adaptive genome refinement did not distort the rank order or relative scale of species-level abundance changes, providing initial quantitative support that warrants further validation across larger spike-in designs and diverse sample matrices.

The DIA results highlight the importance of search-space control. In complex faecal samples, expanding the DIA search space in a single search did not produce a proportional increase in peptide identifications. Comprehensive mode often selected substantially more genomes than Focus mode but produced only modest additional peptide gains, consistent with the sensitivity of current DIA search engines to candidate search-space size. This does not imply that fewer genomes are biologically present in the sample. Rather, it indicates that searching a large candidate genome set all at once can reduce identification depth under current DIA scoring and FDR-control constraints. The iterative Deep strategy addressed this limitation by introducing ranked genomes in smaller batches and accumulating peptide evidence across rounds. This batch-wise strategy allows MetaPilot to explore a broad biologically plausible genome space while avoiding the sensitivity loss associated with a single oversized DIA search.

The DDA-independent reanalysis of the Orbitrap DIA human gut dataset provides the strongest practical demonstration of this principle. DDA-assisted DIA workflows depend on matched DDA data to construct spectral libraries, but such data are not always available in public studies or large cohorts. Without matched DDA data, MetaPilot recovered a deeper peptide and protein evidence space than the DDA-assisted workflow (24.4% more peptides relative to the DDA-derived library; 2.06-fold more than the DDA-assisted DIA search), demonstrating that genome-aware marker evidence can substitute for DDA-derived spectral libraries in cohort-scale reanalysis. The plausibility of the additional identifications was supported by a constrained single-genome validation. When the DIA data were searched against only MGYG000002293 (annotated as *Prevotella sp900557255* in MGnify, corresponding to *Segatella copri*), 40.7% of MetaPilot-exclusive peptides received species-specific Unipept assignments, compared with 25.6% for the peptides shared with metaExpertPro. The fact that the *additional* identifications carry a higher species-specificity than the shared ones argues that they reflect deeper sampling of genuine community members rather than search-space expansion noise, addressing the standard concern that broader search spaces inflate spurious matches.

A related question is whether deeper identification reflects genuine peptide evidence or reduced specificity during adaptive search-space refinement. The defined-mixture analyses provide one orthogonal assessment: despite searching against the full 4,744-genome UHGG catalogue, identifications concentrated on the expected organisms, with approximately 11 of the 12 expected species recovered in the core tier of Qiao_12mix. The cross-biome entrapment analysis extended this assessment to complex faecal samples without defined ground truth. Of 562 tomato-rhizosphere genomes added to the reference catalogue, only one was retained in the DDA refined search space and two in DIA; none reached the DDA core tier and only one reached the DIA core tier. The sole core-tier survivor, *Pseudomonas paraeruginosa*, was a close relative of the strongly supported target-catalogue *P. aeruginosa* representative and was supported by sequence evidence absent from the target-catalogue Pseudomonas genomes. Together, the defined-mixture and cross-biome analyses support the specificity of adaptive genome refinement while highlighting residual ambiguity arising from incomplete strain-level representation.

On the mouse intestinal timsTOF diaPASEF dataset, MetaPilot identified 41.8–119.7% more peptides than uMetaP across the four colonic-content groups. This gain reflects a structural difference between the two workflows: uMetaP depends on a pre-built database expanded with DDA-PASEF-derived *de novo* peptide evidence, whereas MetaPilot constructs its refined search space adaptively from the analysed DIA dataset itself. The same DIA data therefore yielded harmonized peptide, protein, genome, taxonomic, functional and host-associated outputs in a single workflow, with no DDA data required at any stage. MetaPilot’s value, however, extends beyond identification depth. In the Hsp60 epithelial injury model, the injured Δ/ΔIEC_D8 state showed lower peptide and protein identifications, only modest changes in Shannon diversity and a stronger decrease in expressed functional redundancy. At the same time, several core genomes increased rather than decreased. This pattern indicates that the injured D8 metaproteome was not simply a depleted community. Instead, it represented a less redundant expressed ecosystem in which selected genomes became more prominent and carried coherent functional signatures. This example illustrates how a DDA-independent, genome-aware workflow can connect community structure to expressed function within a single dataset — a capability that follows directly from the deeper, harmonized identification space.

Several challenges remain for future development. First, shared peptides among closely related organisms remain an intrinsic source of ambiguity in genome-level interpretation. MetaPilot tracks genome-unique and shared peptide evidence and uses confidence-aware reporting, but no peptide-based metaproteomic workflow can fully remove ambiguity caused by sequence conservation. Second, although MetaPilot reduces dependence on fixed thresholds through greedy genome ranking and knee-based adaptive stopping, some fixed parameters remain in supporting steps. For example, before the greedy search, MetaPilot defines privileged genomes based on genome-unique marker evidence and requires each to contribute at least 1% of the genome-unique peptide pool. This parameter affects early seeding of high-confidence genomes, whereas the main genome selection is determined by marginal peptide gain and adaptive stopping on the ranked genome curve. Future versions should estimate such seeding parameters from dataset size, peptide-evidence distribution and community complexity. Third, the current validation focuses on human gut and mouse intestinal ecosystems. MetaPilot can use the broader set of biome-specific genome catalogues available through MGnify, but some catalogues are substantially larger and more complex than the human gut reference used here.

For example, marine-v2-0 and soil-v1-0 are approximately 2.8-fold and 4.1-fold larger than human-gut-v2-0-2, respectively. These databases may require further optimization of marker filtering, genome-ranking thresholds and adaptive stopping rules to maintain performance across ecosystem scales. Further benchmarking across non-gut biomes, sample matrices, instruments and acquisition designs will therefore be important.

In summary, MetaPilot establishes a unified, genome-aware framework for DDA and DIA metaproteomics. By coupling Marker4 evidence with adaptive genome ranking and refined search-space construction, MetaPilot improves peptide identification depth while preserving genome-resolved taxonomic and functional interpretation. The DDA-independent DIA results show that deep metaproteomic profiling can be achieved without matched DDA-derived libraries, expanding the practical use of public and cohort-scale DIA datasets. As genome catalogues continue to grow across microbial ecosystems, adaptive genome-aware refinement will be increasingly important for scalable and interpretable metaproteomics.

## Methods

### MetaPilot workflow design

MetaPilot was developed as a genome-aware workflow for metaproteomics analysis of DDA and DIA mass spectrometry data. The workflow uses the same conceptual structure for both acquisition modes: spectrum conversion, initial peptide evidence search, genome evidence construction, adaptive genome ranking and selection, refined search-space construction, final identification and quantification, and multi-layer export. In the analyses reported here, DDA data were searched using pFind, and DIA data were searched using DIA-NN. After search-engine parsing, both routes entered the MetaPilot workflow as identified peptide sequences linked to protein identifiers.

The central design of MetaPilot is to use metagenome-assembled genomes as genome-resolved reference units. Each protein sequence in the reference database is linked to its source genome, allowing identified peptides to be mapped from peptide to protein and from protein to genome. Genome-level evidence is then used to select a sample-specific genome set, and proteins from the selected genomes are used to construct the refined search space for final analysis. The final outputs include harmonized peptide, protein, genome, taxonomic and functional tables.

### Reference genome catalogue and Marker4 evidence space

MetaPilot uses genome-resolved databases as reference resources. Each database is organized as per-genome protein FASTA files with associated genome-to-taxonomy metadata and precomputed functional annotation tables. This structure preserves the link between each protein and its source genome throughout the workflow. In this study, the human gut analyses used the MGnify human-gut-v2-0-2 database, corresponding to the Unified Human Gastrointestinal Genome (UHGG) v2.0.2 representative species-level catalogue with 4,744 genomes. The mouse intestinal analysis used the MGnify mouse-gut-v1-0 database with 2,847 representative species-level genomes.

The standard initial evidence search in MetaPilot uses a four-category marker system, referred to as Marker4. Marker4 includes ribosomal proteins, replication/transcription-related proteins, elongation factors and aminoacyl-tRNA synthetases. Marker proteins were extracted from the genome database using protein annotation terms and gene-symbol patterns detailed in **Supplementary Table 5**. The extracted proteins were combined into a single Marker4 search space and used as the standard initial evidence source for both DDA and DIA analyses.

Marker4 extends earlier reduced marker-protein search strategies based on high-abundance or conserved microbial proteins. In our previous MetaLab-related workflow, ribosomal proteins and elongation factors were used as a two-marker evidence space. To evaluate continuity with this earlier strategy, MetaPilot retains a Marker2 mode for DDA analysis using only ribosomal proteins and elongation factors, consistent with earlier marker strategies. Marker4 is the current standard marker strategy in MetaPilot and was used as the main marker design throughout this study.

### Initial peptide evidence searches

For each sample, MetaPilot first performed an initial search against a compact marker-derived search space. Unless otherwise stated, the same enzyme, modification and missed-cleavage settings were used for the corresponding refined search. In DDA workflows, pFind searched the Marker4 or Marker2 database using open-search and semi-digestion settings with a maximum of two missed cleavages. In DIA workflows, DIA-NN searched the Marker4-derived search space using carbamidomethylation of cysteine as a fixed modification; methionine oxidation, protein N-terminal acetylation and protein N-terminal methionine excision as variable settings; tryptic digestion; and a maximum of one missed cleavage. Outputs from both routes were converted into the same MetaPilot evidence type: identified peptide sequences with matched protein identifiers. Search parameters were left at their default values for both engines. pFind results were filtered at 1% FDR at the peptide level and 1% FDR at the protein level, requiring at least one peptide per protein, with precursor mass restricted to 600-100,000 Da and peptide length to 6-100 residues. DIA-NN was run with output filtered at 1% FDR, precursor and protein matrices filtered at 1% precursor- and protein-level FDR, protein matrices additionally filtered at 1% protein q-value, heuristic protein grouping, match-between-runs enabled, a scan window radius of 15, and mass accuracy fixed at 15 ppm for both MS1 and MS2.

For large Marker4 collections, MetaPilot applied an additional search-space reduction step for the initial DIA search. When the Marker4 protein collection contained more than 100,000 sequences, DeepDetect was used to predict peptide detectability from the Marker4 proteins. Peptides with predicted detectability greater than 0.8 were retained to construct a reduced marker peptide database. This reduced database was derived from the same Marker4 evidence space and was used to reduce DIA-NN library generation time and initial DIA search-space burden for large genome catalogues. The Marker4 collections derived from both the human-gut-v2-0-2 and mouse-gut-v1-0 catalogues exceeded this threshold, so the detectability-based reduction was applied in all DIA analyses reported here.

For Seed workflows, the initial search started from user-selected libraries. Identified peptides from these libraries were mapped back to genome-derived proteins using peptide-to-protein mapping tables generated from the reference genome catalogue. After this mapping step, Seed workflows entered the same genome evidence construction and adaptive genome selection procedure as Marker workflows.

For DIA Deep workflows, MetaPilot used the ranked genome list to expand the DIA search space iteratively. Ranked genomes were searched in batches of five genomes per round, and newly identified peptides were accumulated across rounds. Iteration stopped after a minimum of eight rounds when newly added peptides accounted for less than 1.5% of cumulative peptide discoveries in two consecutive rounds. MetaPilot then performed the final DIA-NN quantifying search using the refined search space derived from the accumulated iterative evidence.

### Genome evidence construction

Genome evidence was constructed from peptide identifications generated during the initial marker search. For each identified peptide, MetaPilot recorded the matched protein identifiers, the corresponding genome identifiers and the marker category of the matched protein when applicable. Because a peptide can match proteins from one or multiple genomes, MetaPilot retained the full peptide-to-protein-to-genome mapping rather than assigning each peptide to a single genome at this stage.

Peptides mapping to proteins from only one genome were treated as genome-unique evidence. Peptides mapping to proteins from multiple genomes were retained as shared evidence and contributed to all candidate genomes with explicit ambiguity information. For each candidate genome, MetaPilot calculated the number of identified marker peptides, the number of genome-unique marker peptides, the number of marker categories supported by any identified peptide and the number of marker categories supported by genome-unique peptides. These measurements formed the genome evidence table used for adaptive genome ranking and selection.

### Adaptive genome ranking and selection

Genome selection was performed as a set-cover procedure over the identified marker peptides. Each candidate genome was represented by the set of marker peptides mapped to its proteins. The selected genome set was built to maximize coverage of the observed marker peptide set while limiting redundant genome inclusion.

In the Marker4 workflow, genomes were first filtered by marker-category coverage. The hard filter retained genomes with peptide evidence in the ribosomal, replication/transcription-related and elongation-factor categories. If no genome passed this filter, the filter was relaxed and greedy selection was applied directly to all candidate genomes until the observed marker peptide set was covered.

For genomes passing the marker-category filter, selection proceeded in three stages. First, MetaPilot selected privileged genomes from genomes supported by genome-unique peptides in at least three marker categories. A privileged genome was retained only if its genome-unique peptide count was at least 1% of the total distinct genome-unique peptide set across all candidate genomes. Second, representative genomes from under-covered taxonomic groups were optionally added when they contributed additional marker peptides. Third, all remaining genomes were ordered by greedy marginal gain. At each step, MetaPilot selected the genome adding the largest number of previously uncovered marker peptides. Evidence strength and genome identifier were used to resolve ties in marginal gain.

For each greedy step, MetaPilot recorded the number of newly covered marker peptides, cumulative peptide coverage and marginal-gain ratio. The marginal-gain ratio was smoothed along the ranked genome list and used to identify stopping points. Because Marker2 is retained only for DDA, DDA modes are named for the marker set used (Marker2- or Marker4-), whereas DIA modes, which always use Marker4, are named simply Marker-; modes are therefore written throughout as Marker4-Focus, Marker4-Comprehensive, Marker-Focus and Marker-Deep. Focus and Comprehensive modes used the same ranked genome list but different stopping rules. Focus stopped at the first knee of the genome-coverage curve, yielding the most compact refined search space, whereas Comprehensive continued genome addition beyond the first knee to maximize candidate coverage in a single round. In DIA Deep mode, ranked genomes were introduced in successive batches and peptide evidence was accumulated iteratively until a plateau stopping criterion was met, before the final quantifying search.

### Refined search-space construction and final quantification

After genome selection, MetaPilot constructed a refined sample-specific microbial search space by concatenating all proteins from the selected genomes. For DDA workflows, host proteins were appended to the microbial FASTA when host analysis was enabled, and the combined microbial-host database was searched for final identification and quantification.

For DIA Marker and iterative Seed workflows, proteins from selected genomes were used to build the refined microbiome DIA search space for the final DIA-NN quantifying search. In mixed Seed workflows, the refined microbiome search space was converted into the required library format when it needed to be combined with standalone user-selected libraries. When host analysis was enabled, the host database was searched separately at q ≤ 0.01 at both the precursor and protein levels. Microbial and host results were exported as separate peptide- and protein-level tables and integrated into the MetaPilot output structure.

In DIA Deep mode, ranked genomes were searched iteratively in batches before the final quantifying search. Peptides identified across iterative rounds were accumulated until the stopping criterion was reached. The accumulated evidence was then used to construct the final refined DIA search space, and the resulting peptide- and protein-level matrices were converted into MetaPilot’s harmonized output format.

### Genome and protein confidence scoring

After final identification, MetaPilot computed genome- and protein-level confidence scores from peptide evidence. Peptide evidence was collapsed at the sequence level and mapped to the proteins and genomes through protein identifiers. Genome-level confidence scoring was performed independently of the marker-based genome-selection step.

For genome scoring, only genome-unique peptides were used. Each peptide was assigned a weighted evidence score combining identification confidence and quantitative intensity. For genome *g*, the genome score *S_g_* was calculated as the sum of the five highest weighted peptide scores assigned to that genome:

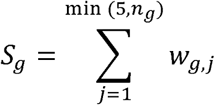

where *w_g,i_* denotes the ranked weighted evidence score of genome-unique peptide *j* for genome *g*, and *n_g_* is the number of genome-unique peptides assigned to that genome.

Empirical genome-level significance was estimated by decoy resampling. For each genome, size-matched sets of decoy peptide scores were sampled 10,000 times. The empirical p-value was calculated as:

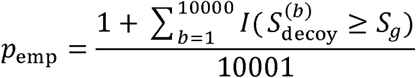

where 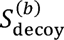 is the top-five summed score from the *b*-th decoy resampling. Empirical p-values were adjusted across genomes using the Benjamini–Hochberg procedure.

For protein scoring, all weighted peptide evidence assigned to each protein was summed. Protein-level empirical p-values and q-values were estimated using the same decoy-resampling and multiple-testing correction procedure. This scoring step supported confidence-aware genome and protein reporting, particularly when final identifications included peptides shared among closely related genomes.

### Genome tiering, taxonomic reporting and functional reporting

In addition to statistical scoring, MetaPilot assigned each reported genome to a qualitative tier derived from the genome-ranking procedure used during search-space construction. During greedy ranking, each genome was ordered according to the number of previously uncovered marker peptides it contributed. For greedy step *i*, MetaPilot recorded the incremental gain *g_i_*, the cumulative number of covered marker peptides *C_i_*, and the incremental contribution ratio:

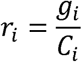

The ratio series was smoothed using a five-step moving average:

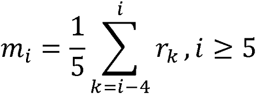

Knee positions were estimated from the smoothed curve using chord-distance, kneedle-style, curvature-based, piecewise-linear and slope-change criteria. The Focus threshold was defined as the median of the kneedle-style, curvature and slope-change knees. The Comprehensive threshold was defined as the later of the chord-distance and piecewise-linear knees. Genomes ranked before or at the Focus threshold were labelled core, genomes between the Focus and Comprehensive thresholds were labelled mid, and later contributing genomes were labelled tail. Genomes detected in the final abundance table but not assigned during the ranking step were also labelled tail.

Taxonomic reports were generated by linking genome identifiers to the taxonomy metadata distributed with the MetaPilot genome database. They are downloaded as database packages containing per-genome protein FASTA files, genome-to-lineage tables and precomputed functional annotation tables. Genome-level peptide counts, unique peptide counts, quantitative intensities, empirical confidence scores and reporting tiers were first summarized at genome level and then aggregated across standard taxonomic ranks. GTDB (Genome Taxonomy Database, https://gtdb.ecogenomic.org/) lineage information was used as the primary genome taxonomy, and NCBI lineage information was included when available through database mapping.

Functional reporting was performed by mapping identified genome proteins to the precomputed functional annotation tables included in the corresponding genome database package. For the MGnify-derived databases used in this study, functional annotations were based on the catalogue-provided eggNOG-style annotation results. Version-specific annotation details for each catalogue are documented in the MGnify Genomes database release notes (https://docs.mgnify.org/). Protein evidence was aggregated to functional terms to generate summaries for Gene Ontology terms, Enzyme Commission identifiers, KEGG orthologs, BRITE terms, COG accessions, NOG accessions and COG/NOG functional categories. These functional layers were then used to generate quantitative matrices for downstream comparison of expressed microbial functions across samples and groups.

### Evaluation datasets and comparative analyses

#### Paired DDA and DIA benchmark datasets

MetaPilot was evaluated using paired DDA and DIA metaproteomics datasets from the Elo and Qiao studies. These datasets included defined microbial mixtures and complex faecal samples and were used to assess genome selection, peptide identification and comparison with published workflows across sample complexity and acquisition mode.

For peptide-level comparisons, representative MetaPilot outputs were selected from the workflow modes used in the Results: Marker4-Focus for simple DDA datasets, Marker4-Comprehensive for faecal DDA datasets, Marker-Focus for simple DIA datasets and Marker-Deep for faecal DIA datasets. Peptide sequences from MetaPilot and published result tables were standardized by removing non-letter modification or flanking-residue annotations, converting sequences to uppercase and treating isoleucine and leucine as equivalent by mapping I to L. Decoy and contaminant entries were removed before comparison. Peptide-set overlap was visualized using Venn diagrams or UpSet plots depending on the number of compared workflows. For the Qiao spike-in faecal benchmark, MetaPilot-derived species abundance ratios were compared with the expected spike-in ratios.

#### Entrapment analysis

To assess whether adaptive refinement concentrates identifications on genuinely present organisms in samples of unknown composition — complementing the known-composition validation from the 12-species mixture — we constructed an entrapment database from a taxonomically distinct tomato-rhizosphere genome catalogue. Candidate entrapment genomes sharing species-level GTDB taxonomy with any genome in the 4,744-genome UHGG target catalogue were excluded (17 genomes removed) to reduce direct taxonomic overlap between the target and entrapment catalogues; the remaining 562 genomes were merged with the target catalogue to form a combined 5,306-genome search database. The entrapment set was sized at approximately 12% of the target catalogue by design, to limit the increase in search runtime relative to the target-only catalogue used elsewhere in this study.

The combined catalogue was searched against Qiao_faecal_DDA and Qiao_faecal_DIA using Marker4-Comprehensive and Marker-Deep, respectively, with the same parameters used for the corresponding target-only analyses. Genomes were evaluated at three stages: the candidate pool, comprising genomes passing an initial genome-level empirical significance test (p_emp and Benjamini– Hochberg-adjusted q-value); the final refined search space, comprising genomes retained after greedy/knee-based selection; and the core tier, the highest-confidence subset of the final refined search space assigned by the same selection procedure (as distinct from mid- and tail-confidence genomes).

#### DDA-independent reanalysis of the metaExpertPro DIA dataset

MetaPilot was applied to the Orbitrap Exploris 480 DIA data from the published metaExpertPro study in DDA-independent Marker-Deep mode. The analysis used Marker4 evidence, genome-aware ranking, iterative refinement and final DIA-NN quantification without using the matched DDA-derived spectral library from the original workflow. The host search used a human UniProt protein database downloaded on 1 September 2025, containing 20,432 proteins.

MetaPilot peptide and protein identifications were compared with the published metaExpertPro DDA-derived spectral library and DIA identification results. A constrained single-genome validation was performed using MGYG000002293, annotated as *Prevotella sp900557255* in MGnify and corresponding to *Segatella copri* under current taxonomy. DIA data were searched against this single genome, and the resulting peptide evidence was compared with the MetaPilot and metaExpertPro peptide sets. Unipept analyses used web application version 6.5.0 with UniProtKB_2025.04.

#### DDA-independent reanalysis of the uMetaP DIA dataset

MetaPilot was applied to the published mouse intestinal colonic-content DIA dataset originally analysed using uMetaP. MetaPilot was run in DDA-independent DIA mode using the MGnify mouse-gut-v1-0 genome catalogue, containing 2,847 representative species-level genomes in the local MetaPilot database configuration. The host search used a mouse UniProt protein database downloaded on 5 May 2026, containing 17,266 proteins.

Principal component analysis was performed at genome and annotated-protein levels. Shannon diversity was calculated from genome-level abundance profiles. Pairwise comparisons shown for Shannon diversity and expressed functional redundancy were performed using two-sided Mann–Whitney U tests. Because these comparisons involve small group sizes, the Mann-Whitney U statistic takes discrete values and identical p-values can arise for independent comparisons that yield the same rank configuration.

For altered-feature analysis, each feature was compared between predefined group contrasts using a two-sided Mann–Whitney U test. P-values were adjusted within each analytical layer using the Benjamini–Hochberg procedure. Features were considered altered when BH-adjusted FDR < 0.01 and |log2 fold change| ≥ 1. Altered-feature rates were calculated as the proportion of altered features among all tested features within each analytical layer.

Expressed functional redundancy was calculated as the effective number of genome contributors per KEGG KO. For each sample *s*, KEGG KO *k*, and contributing genome *g*, the genome contribution fraction was calculated as:

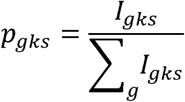

where *I_gks_* is the abundance contribution of genome*g* to KO *k* in sample *s*. The effective number of contributing genomes was then calculated as:

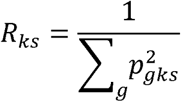

This metric equals one when a KO is supported by a single genome exclusively, and approaches the total number of contributing genomes as signal becomes evenly distributed. Differential genome analysis was used to identify altered genomes across group comparisons, and increased core genomes were linked to shared KEGG KO trends for genome-to-function interpretation.

## Data availability

No new human or animal data were generated. All datasets were obtained from public repositories under the accessions listed below. The Elo_12mix and Elo_faecal datasets were downloaded from PRIDE under accession PXD008738. The Qiao_12mix and Qiao_faecal datasets were downloaded from iProX under accession IPX0003851000. The metaExpertPro Guangzhou Nutrition and Health Study dataset was downloaded from PRIDE under accession PXD051104. The uMetaP colonic-content dataset was downloaded from PRIDE under accession PXD051792.

## Code availability

MetaPilot is implemented as a software platform integrating pFind, DIA-NN, genome selection, post-processing and export modules. Analyses in this manuscript were performed using MetaPilot 1.0.0-beta.6, DIA-NN 2.3.0, pFind 3.2.3 and DeepDetect 1.0.0. Workflow parameters were exported as JSON configuration files, and selected genome lists and output tables were retained for reproducibility. Source code, documentation and the GUI and HPC-compatible command-line workflows are publicly available at https://github.com/cksakura/MetaPilot under the Apache-2.0 licence. All releases, including version 1.0.0-beta.6 used for the analyses reported here and the current stable release 1.1.0, are archived at https://github.com/cksakura/MetaPilot-Releases/releases.

## Acknowledgements

The authors gratefully acknowledge the support of the Biotechnology and Biological Sciences Research Council (BBSRC); this research was funded by the BBSRC Institute Strategic Programme Food Microbiome and Health BB/X011054/1 and its constituent project(s) BBS/E/QU/230001B.

## Author contributions

K.C. conceived the study, designed and implemented the MetaPilot workflow, performed all data processing, benchmarking and downstream analyses, generated the figures, interpreted the results and wrote the manuscript. D.F. provided scientific supervision, contributed to study design and interpretation, and critically revised the manuscript. Both authors discussed the results, contributed to manuscript editing and approved the final version.

## References

1. Kleiner, M. Metaproteomics: Much More than Measuring Gene Expression in Microbial Communities. mSystems 4 (2019).

2. Tanca, A. et al. The impact of sequence database choice on metaproteomic results in gut microbiota studies. Microbiome 4, 51 (2016).

3. Heyer, R. et al. Challenges and perspectives of metaproteomic data analysis. J Biotechnol 261, 24–36 (2017).

4. Li, J. et al. An integrated catalog of reference genes in the human gut microbiome. Nat Biotechnol 32, 834–841 (2014).

5. Chatterjee, S. et al. A comprehensive and scalable database search system for metaproteomics. BMC Genomics 17, 642 (2016).

6. Rajczewski, A.T. et al. Data-Independent Acquisition Mass Spectrometry as a Tool for Metaproteomics: Interlaboratory Comparison Using a Model Microbiome. Proteomics 25, e202400187 (2025).

7. Aakko, J. et al. Data-Independent Acquisition Mass Spectrometry in Metaproteomics of Gut Microbiota-Implementation and Computational Analysis. J Proteome Res 19, 432–436 (2020).

8. Pietila, S., Suomi, T. & Elo, L.L. Introducing untargeted data-independent acquisition for metaproteomics of complex microbial samples. ISME Commun 2, 51 (2022).

9. Almeida, A. et al. A unified catalog of 204,938 reference genomes from the human gut microbiome. Nat Biotechnol 39, 105–114 (2021).

10. Gurbich, T.A. et al. MGnify Genomes: A Resource for Biome-specific Microbial Genome Catalogues. J Mol Biol 435, 168016 (2023).

11. Meier, F. et al. diaPASEF: parallel accumulation-serial fragmentation combined with data-independent acquisition. Nat Methods 17, 1229–1236 (2020).

12. Zhang, X. et al. MetaPro-IQ: a universal metaproteomic approach to studying human and mouse gut microbiota. Microbiome 4, 31 (2016).

13. Cheng, K. et al. MetaLab: an automated pipeline for metaproteomic data analysis. Microbiome 5 (2017).

14. Cheng, K. et al. MetaLab 2.0 Enables Accurate Post-Translational Modifications Profiling in Metaproteomics. J Am Soc Mass Spectr 31, 1473– 1482 (2020).

15. Cheng, K. et al. MetaLab-MAG: A Metaproteomic Data Analysis Platform for Genome-Level Characterization of Microbiomes from the Metagenome-Assembled Genomes Database. J Proteome Res 22, 387–398 (2023).

16. Stamboulian, M., Li, S. & Ye, Y. Using high-abundance proteins as guides for fast and effective peptide/protein identification from human gut metaproteomic data. Microbiome 9, 80 (2021).

17. Yang, J., Cheng, Z., Gong, F. & Fu, Y. DeepDetect: Deep Learning of Peptide Detectability Enhanced by Peptide Digestibility and Its Application to DIA Library Reduction. Anal Chem 95, 6235–6243 (2023).

18. Zhao, J. et al. Data-independent acquisition boosts quantitative metaproteomics for deep characterization of gut microbiota. NPJ Biofilms Microbiomes 9, 4 (2023).

19. Wu, E. et al. High-Abundance Protein-Guided Hybrid Spectral Library for Data-Independent Acquisition Metaproteomics. Anal Chem 96, 1029–1037 (2024).

20. Mesuere, B. et al. Unipept: tryptic peptide-based biodiversity analysis of metaproteome samples. J Proteome Res 11, 5773–5780 (2012).

21. Gurdeep Singh, R., et al. Unipept 4.0: Functional Analysis of Metaproteome Data. J Proteome Res 18, 606–615 (2019).

22. Wicaksono, W.A. et al. The edible plant microbiome: evidence for the occurrence of fruit and vegetable bacteria in the human gut. Gut Microbes 15, 2258565 (2023).

23. Turnbull, P.C. & Kramer, J.M. Intestinal carriage of Bacillus cereus: faecal isolation studies in three population groups. J Hyg (Lond) 95, 629–638 (1985).

24. Ohara, T. & Itoh, K. Significance of Pseudomonas aeruginosa colonization of the gastrointestinal tract. Intern Med 42, 1072–1076 (2003).

25. Ting, Y.S. et al. PECAN: library-free peptide detection for data-independent acquisition tandem mass spectrometry data. Nat Methods 14, 903–908 (2017).

26. Bruderer, R. et al. Extending the limits of quantitative proteome profiling with data-independent acquisition and application to acetaminophen-treated three-dimensional liver microtissues. Mol Cell Proteomics 14, 1400–1410 (2015).

27. Demichev, V., Messner, C.B., Vernardis, S.I., Lilley, K.S. & Ralser, M. DIA-NN: neural networks and interference correction enable deep proteome coverage in high throughput. Nat Methods 17, 41–44 (2020).

28. Sun, Y. et al. metaExpertPro: A Computational Workflow for Metaproteomics Spectral Library Construction and Data-Independent Acquisition Mass Spectrometry Data Analysis. Mol Cell Proteomics 23, 100840 (2024).

29. Xian, F. et al. Ultra-sensitive metaproteomics redefines the dark metaproteome, uncovering host-microbiome interactions and drug targets in intestinal diseases. Nat Commun 16, 6644 (2025).

