## Supplementary Information for "MetaPilot enables adaptive genome-aware DDA and DIA metaproteomics using microbial genome catalogues"

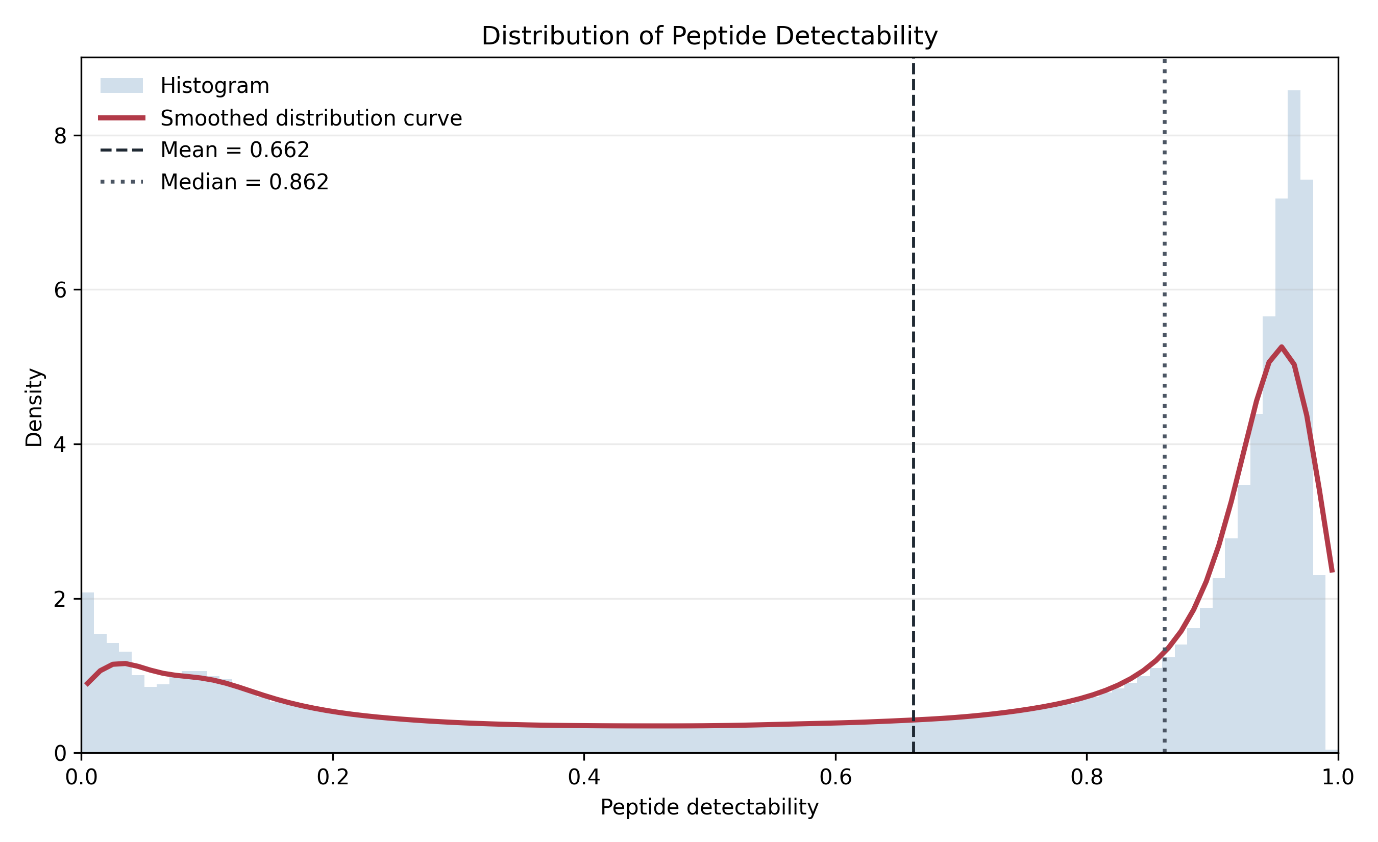

**Figure S1.** **Distribution of predicted peptide detectability scores for the peptide list collected from 15 metaproteomics projects.** Peptide detectability was predicted for each peptide by DeepDetect, where higher values indicate a greater likelihood of detection by mass spectrometry. The distribution shows that a large proportion of peptides have high predicted detectability, with a median detectability of 0.862. For the DIA initial search, peptides with detectability scores above 0.8 were selected to reduce the search space while retaining peptides most likely to be observed experimentally. This cutoff is therefore a reasonable strategy for improving search efficiency without broadly excluding the highly detectable peptide population.

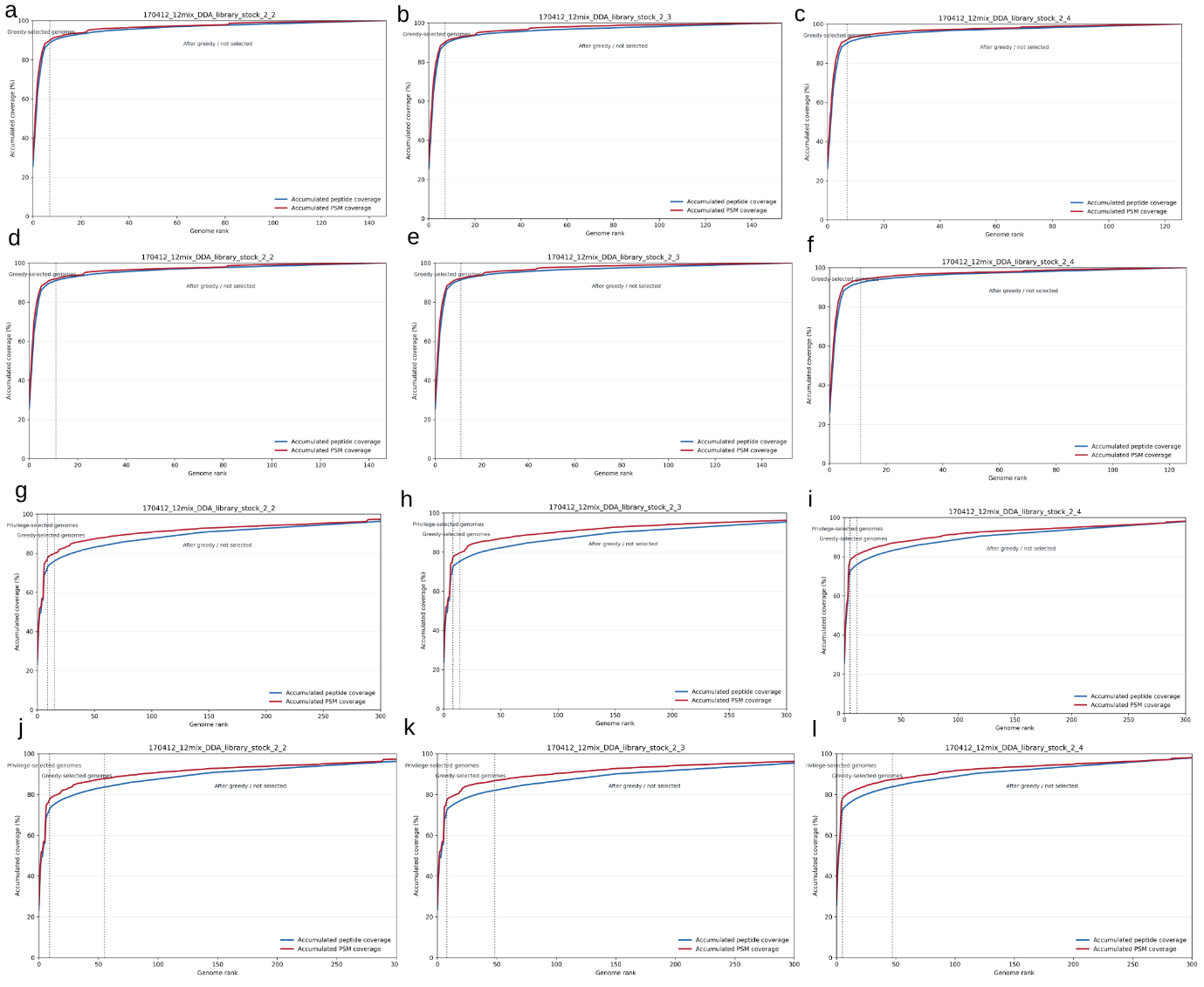

**Figure S2. Marker-peptide coverage and genome selection behaviour in Elo_12mix_DDA samples.** For Figures S2-S16, accumulated marker-peptide and PSM coverage curves were used to evaluate how MetaPilot ranks and selects genomes for refined database construction. Blue lines indicate accumulated marker peptide coverage, and red lines indicate accumulated PSM coverage. Vertical dashed lines mark the genome-selection boundaries used by MetaPilot, including privileged-selected genomes where present and the greedy-selected genome cutoff. In this figure columns correspond to the three raw files, and rows correspond to four DDA genome-selection modes: Marker2 Focus, Marker2 Comprehensive, Marker4 Focus and Marker4 Comprehensive. In the Marker2 workflows, coverage reached a plateau rapidly, indicating that a small number of ranked genomes explained most of the marker evidence in the defined 12-species mixture. Marker4 workflows selected broader genome sets, particularly in Comprehensive mode, while maintaining early high coverage of the observed marker peptide and PSM evidence. This supports the use of adaptive genome selection to construct compact refined databases for low-complexity samples.

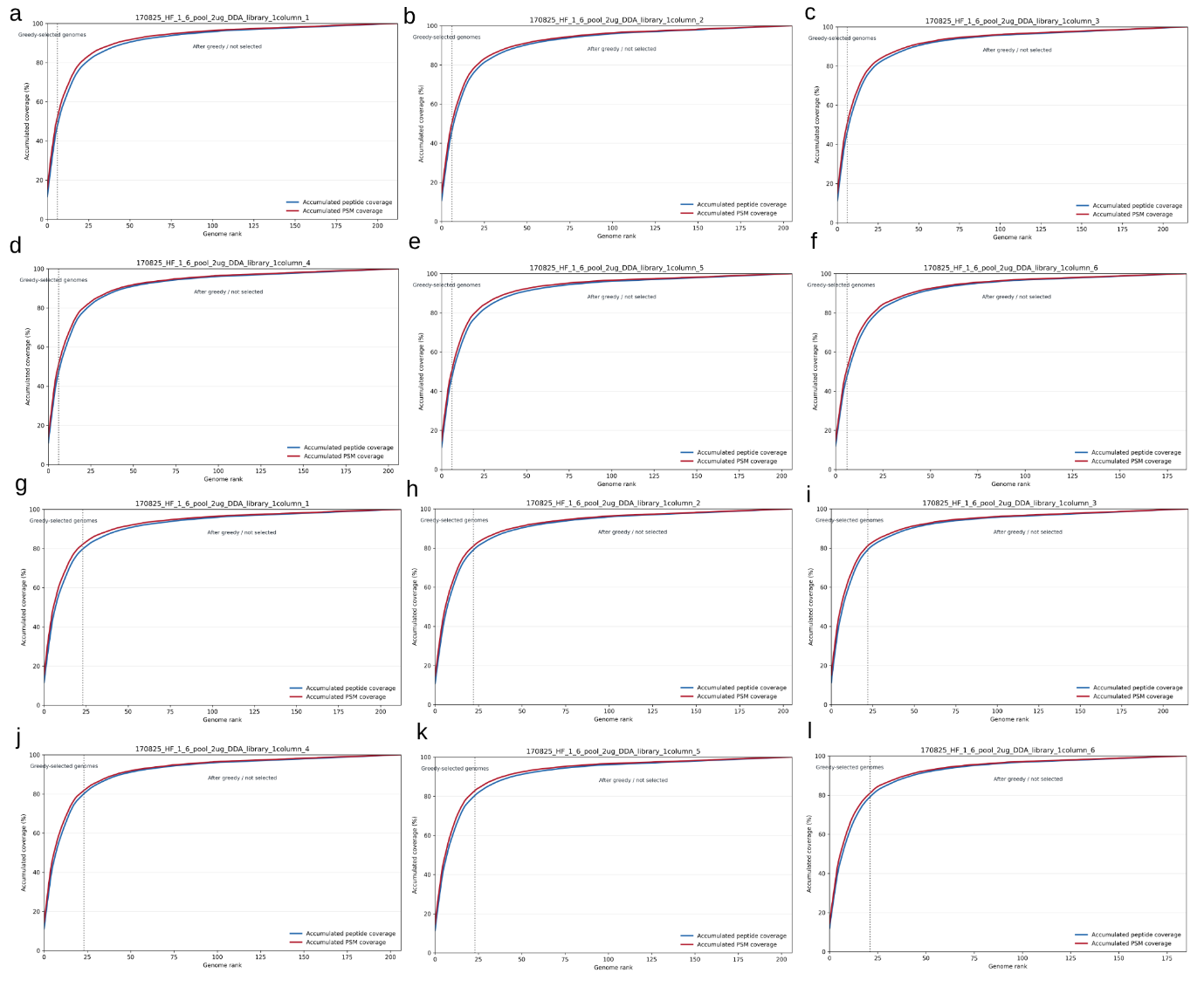

**Figure S3. Marker-peptide coverage and genome selection behaviour in Elo_fecal_DDA samples by the Marker2 strategy.** This dataset contains 6 raw files, a-f represent the Marker2 Focus strategy and g-l represent the Marker2 Comprehensive strategy. Each DDA raw file is shown separately because stochastic precursor selection can lead to run-specific peptide evidence and genome-selection behaviour.

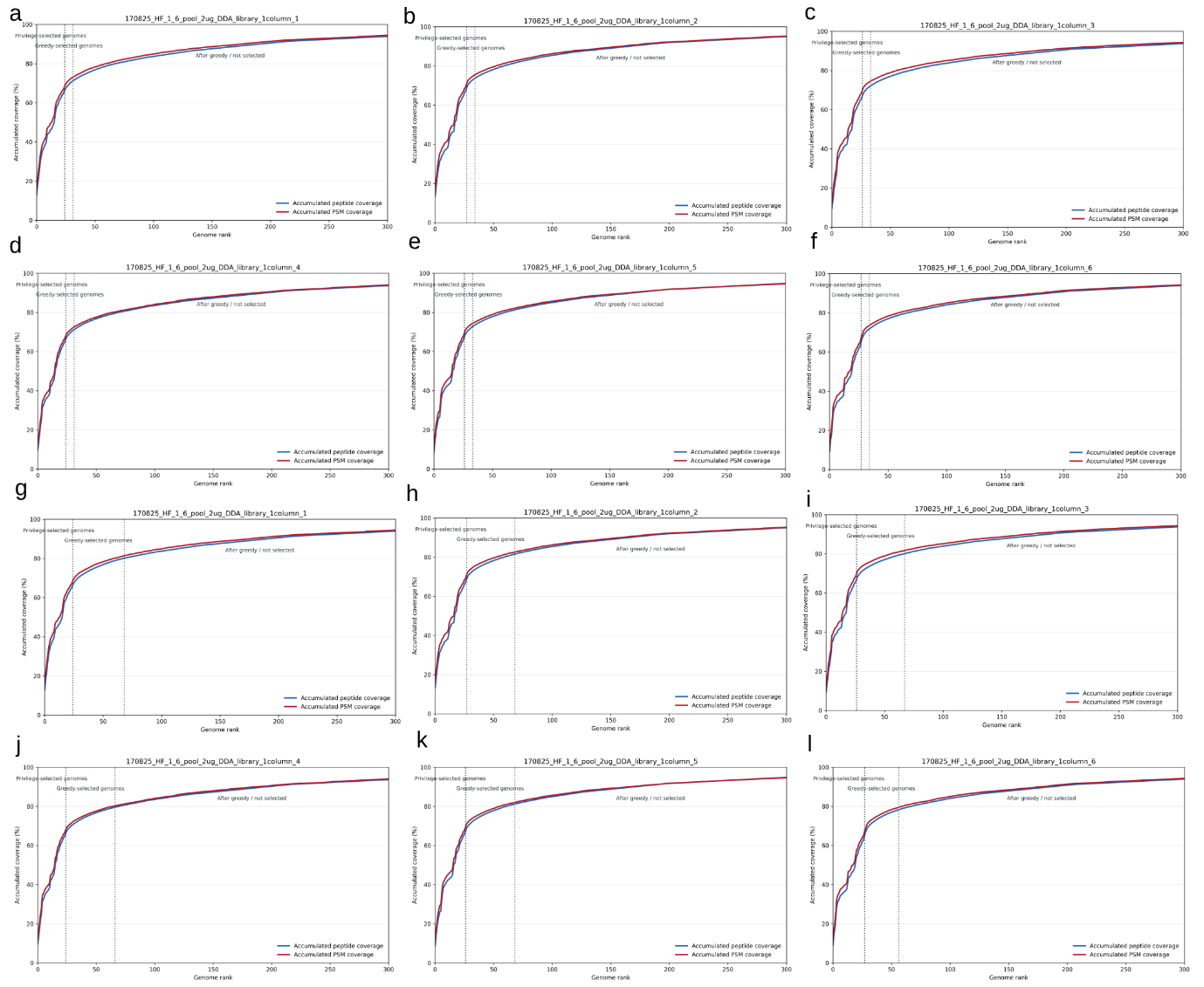

**Figure S4. Marker-peptide coverage and genome selection behaviour in Elo_fecal_DDA samples by the Marker4 strategy.** This dataset contains 6 raw files, a-f represent the Marker4 Focus strategy and g-l represent the Marker4 Comprehensive strategy. Each DDA raw file is shown separately because stochastic precursor selection can lead to run-specific peptide evidence and genome-selection behaviour.

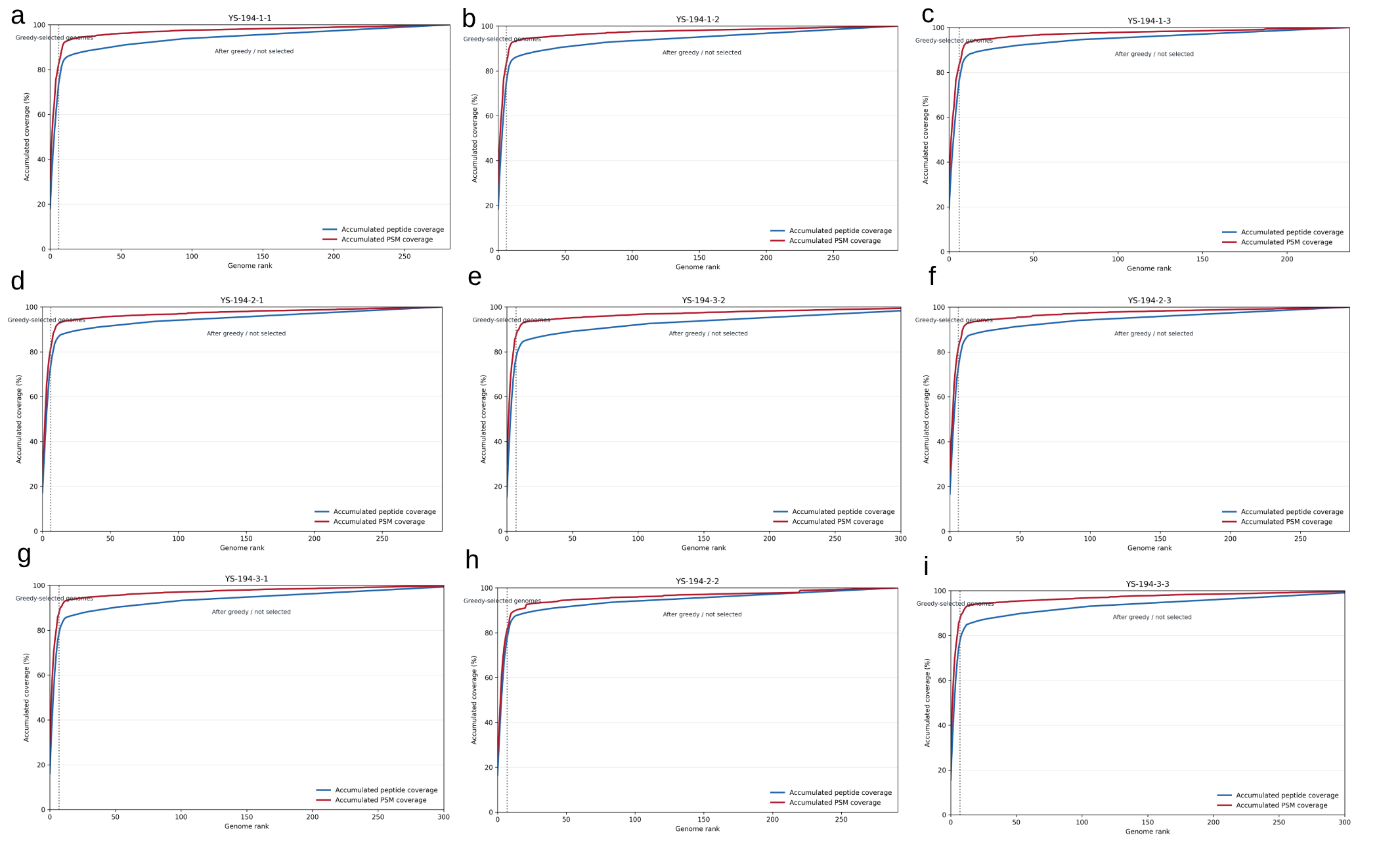

**Figure S5. Marker-peptide coverage and genome selection behaviour in Qiao_12mix_DDA samples by the Marker2 focus strategy.** This dataset contains 9 raw files. Each DDA raw file is shown separately because stochastic precursor selection can lead to run-specific peptide evidence and genome-selection behaviour.

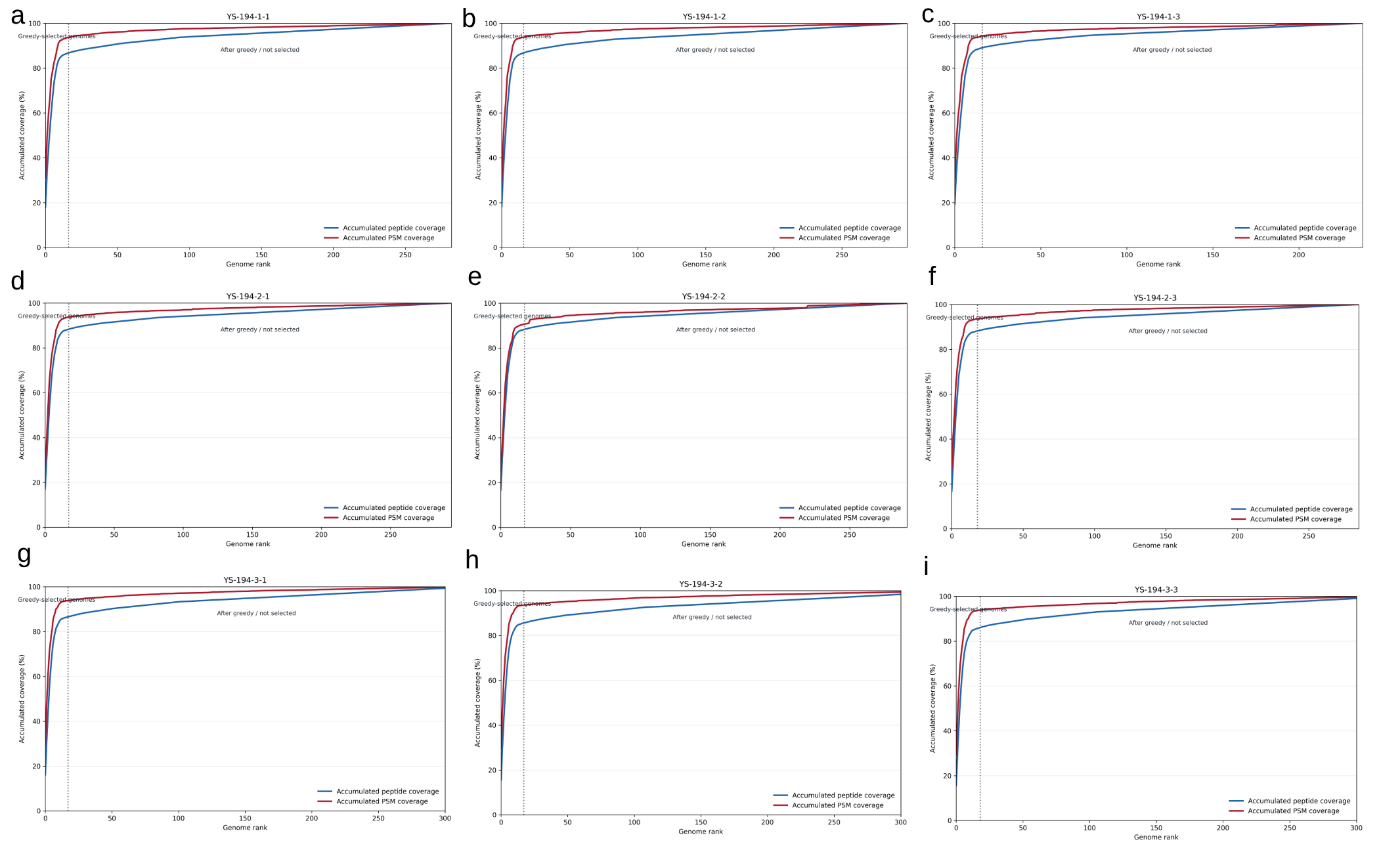

**Figure S6. Marker-peptide coverage and genome selection behaviour in Qiao_12mix_DDA samples by the Marker2 comprehensive strategy.** This dataset contains 9 raw files. Each DDA raw file is shown separately because stochastic precursor selection can lead to run-specific peptide evidence and genome-selection behaviour.

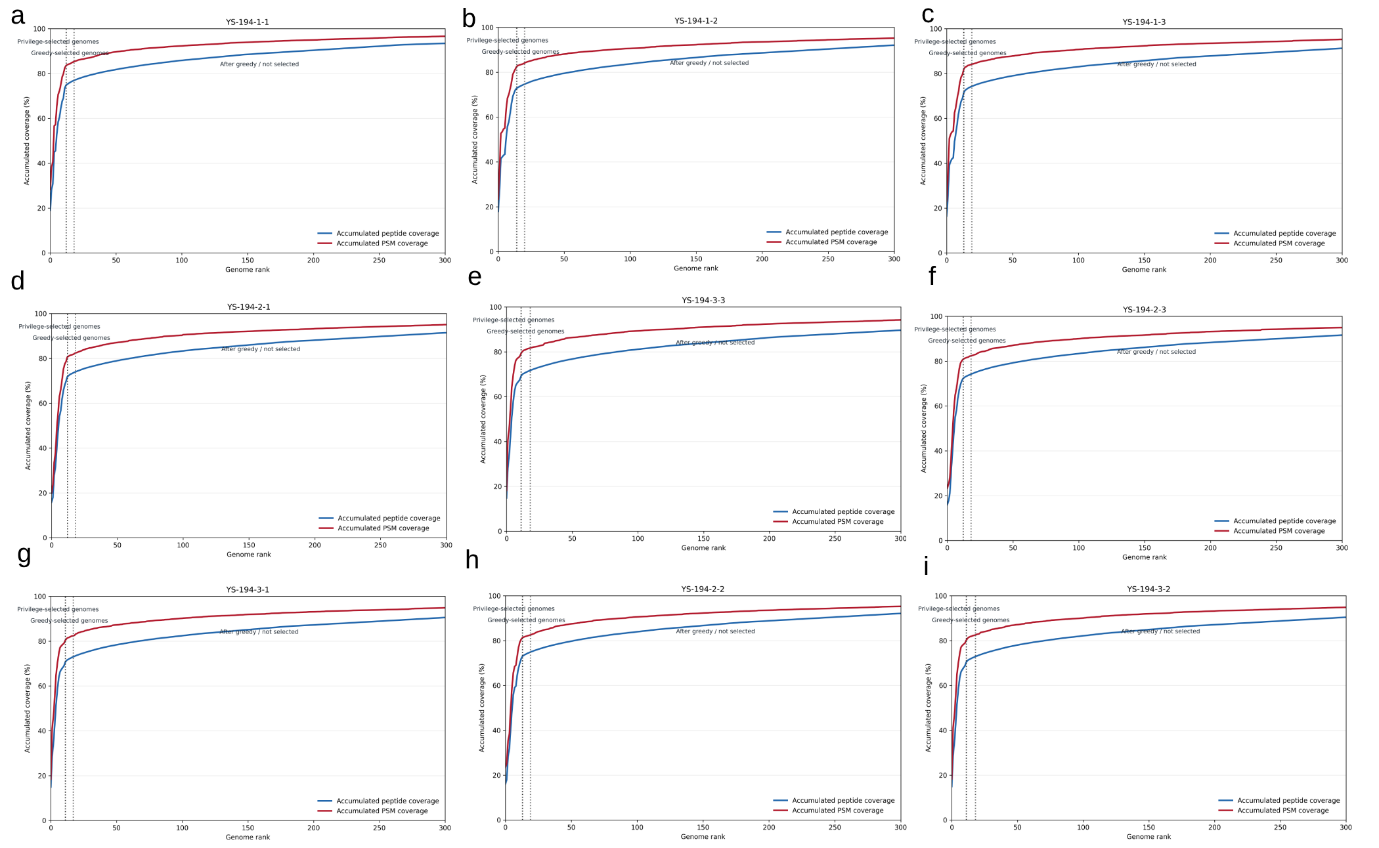

**Figure S7. Marker-peptide coverage and genome selection behaviour in Qiao_12mix_DDA samples by the Marker4 focus strategy.** This dataset contains 9 raw files. Each DDA raw file is shown separately because stochastic precursor selection can lead to run-specific peptide evidence and genome-selection behaviour.

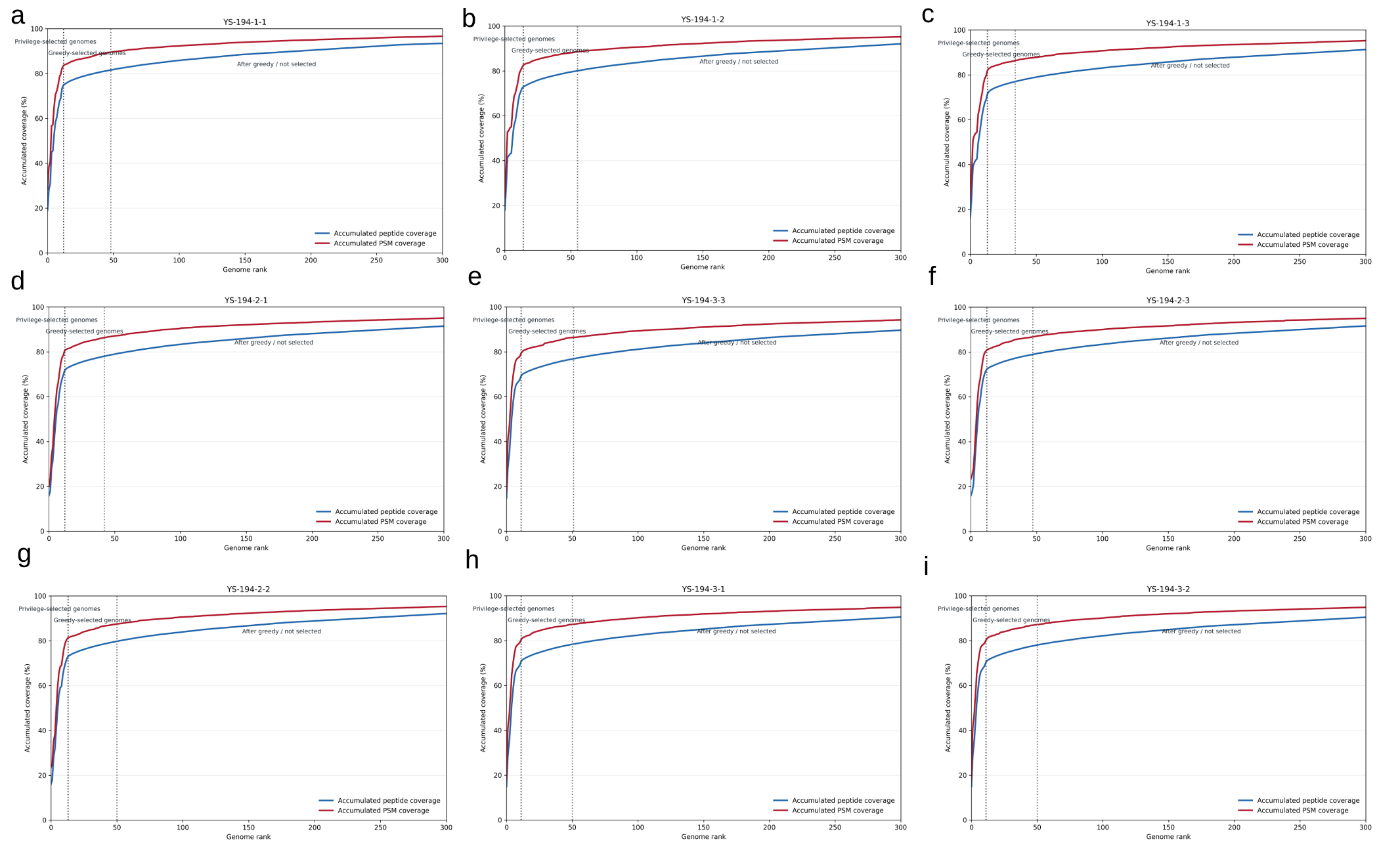

**Figure S8. Marker-peptide coverage and genome selection behaviour in Qiao_12mix_DDA samples by the Marker4 comprehensive strategy.** This dataset contains 9 raw files. Each DDA raw file is shown separately because stochastic precursor selection can lead to run-specific peptide evidence and genome-selection behaviour.

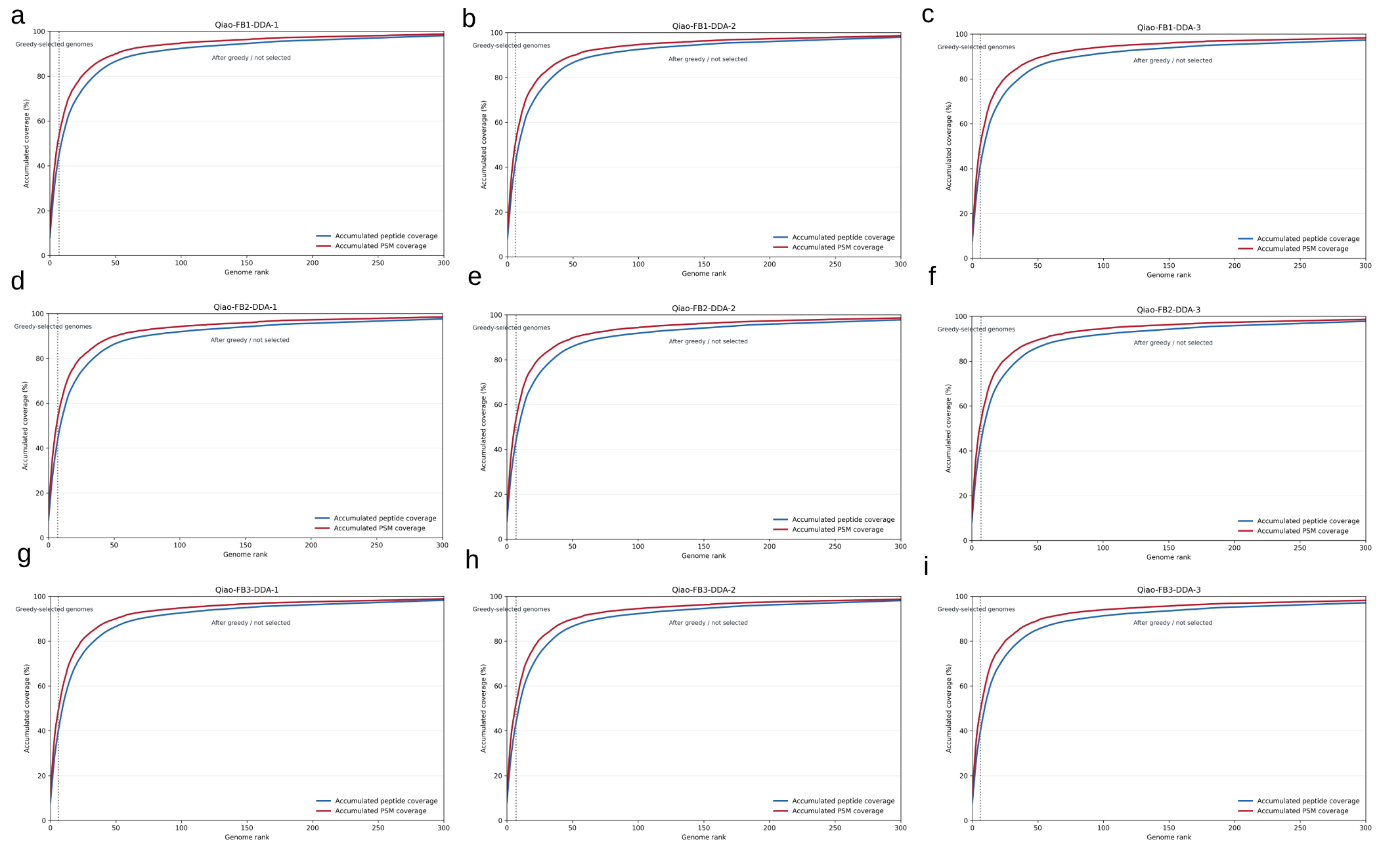

**Figure S9. Marker-peptide coverage and genome selection behaviour in Qiao_fecal_DDA samples by the Marker2 focus strategy.** This dataset contains 9 raw files. Each DDA raw file is shown separately because stochastic precursor selection can lead to run-specific peptide evidence and genome-selection behaviour.

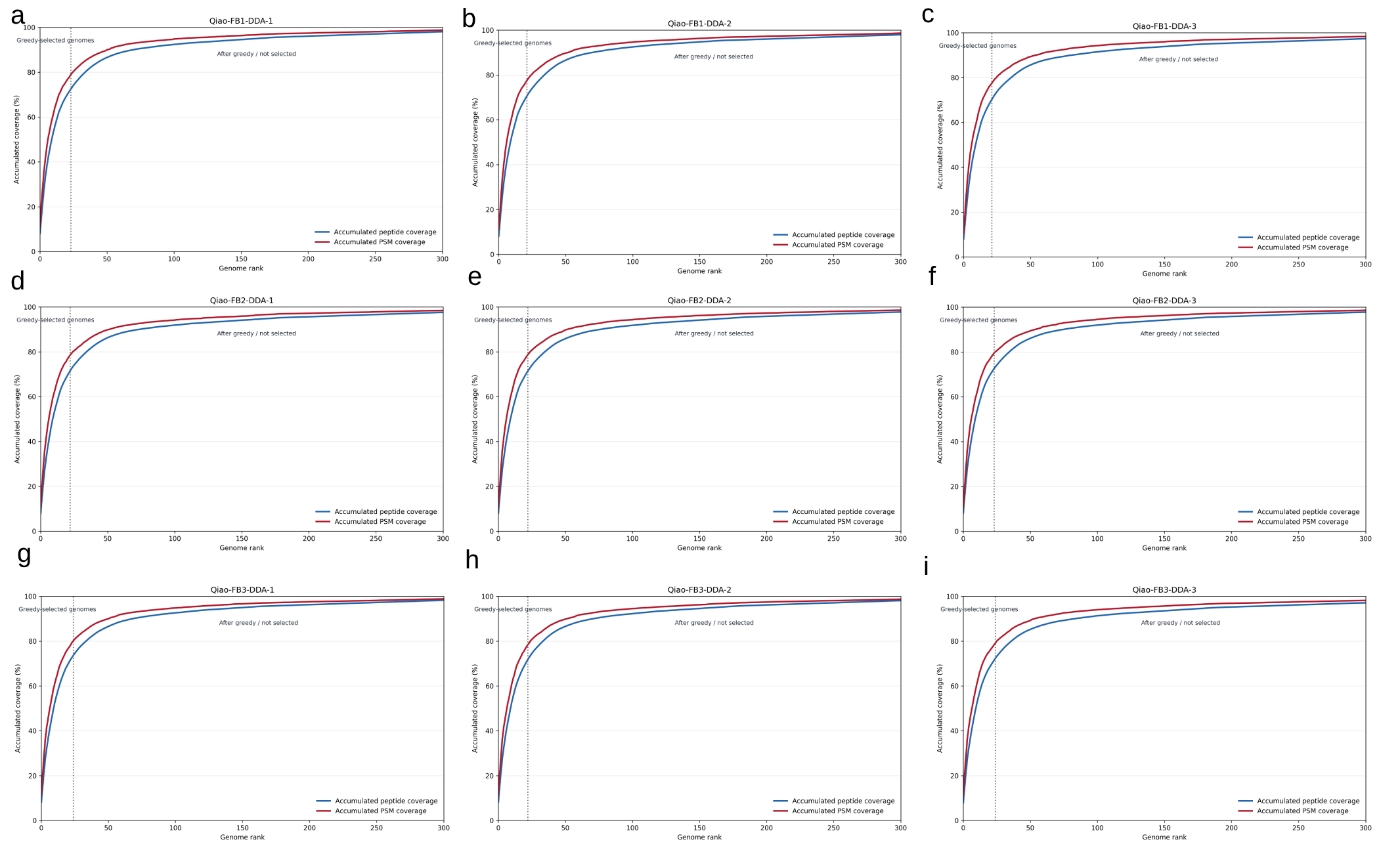

**Figure S10. Marker-peptide coverage and genome selection behaviour in Qiao_fecal_DDA samples by the Marker2 comprehensive strategy.** This dataset contains 9 raw files. Each DDA raw file is shown separately because stochastic precursor selection can lead to run-specific peptide evidence and genome-selection behaviour.

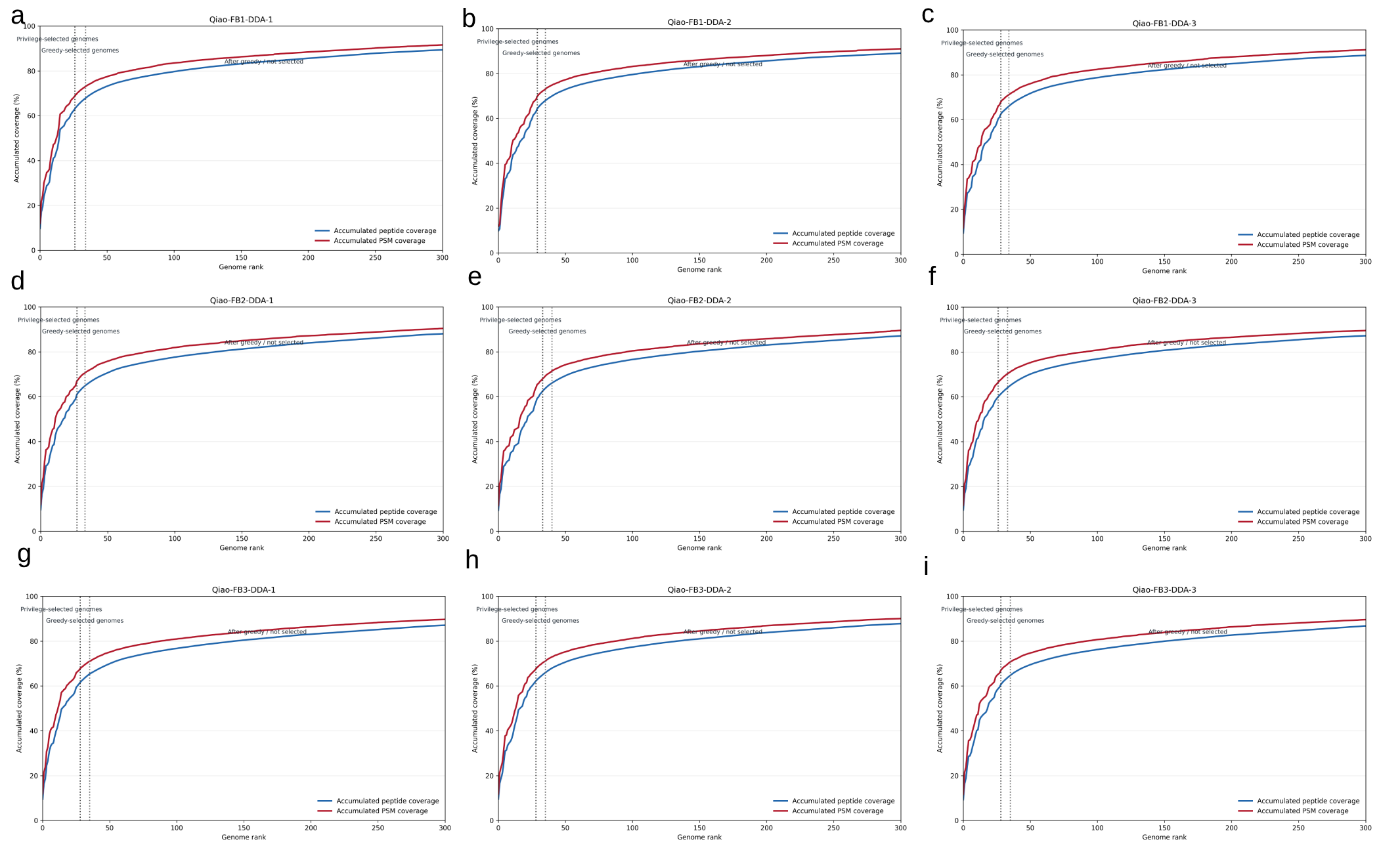

**Figure S11. Marker-peptide coverage and genome selection behaviour in Qiao_fecal_DDA samples by the Marker4 focus strategy.** This dataset contains 9 raw files. Each DDA raw file is shown separately because stochastic precursor selection can lead to run-specific peptide evidence and genome-selection behaviour.

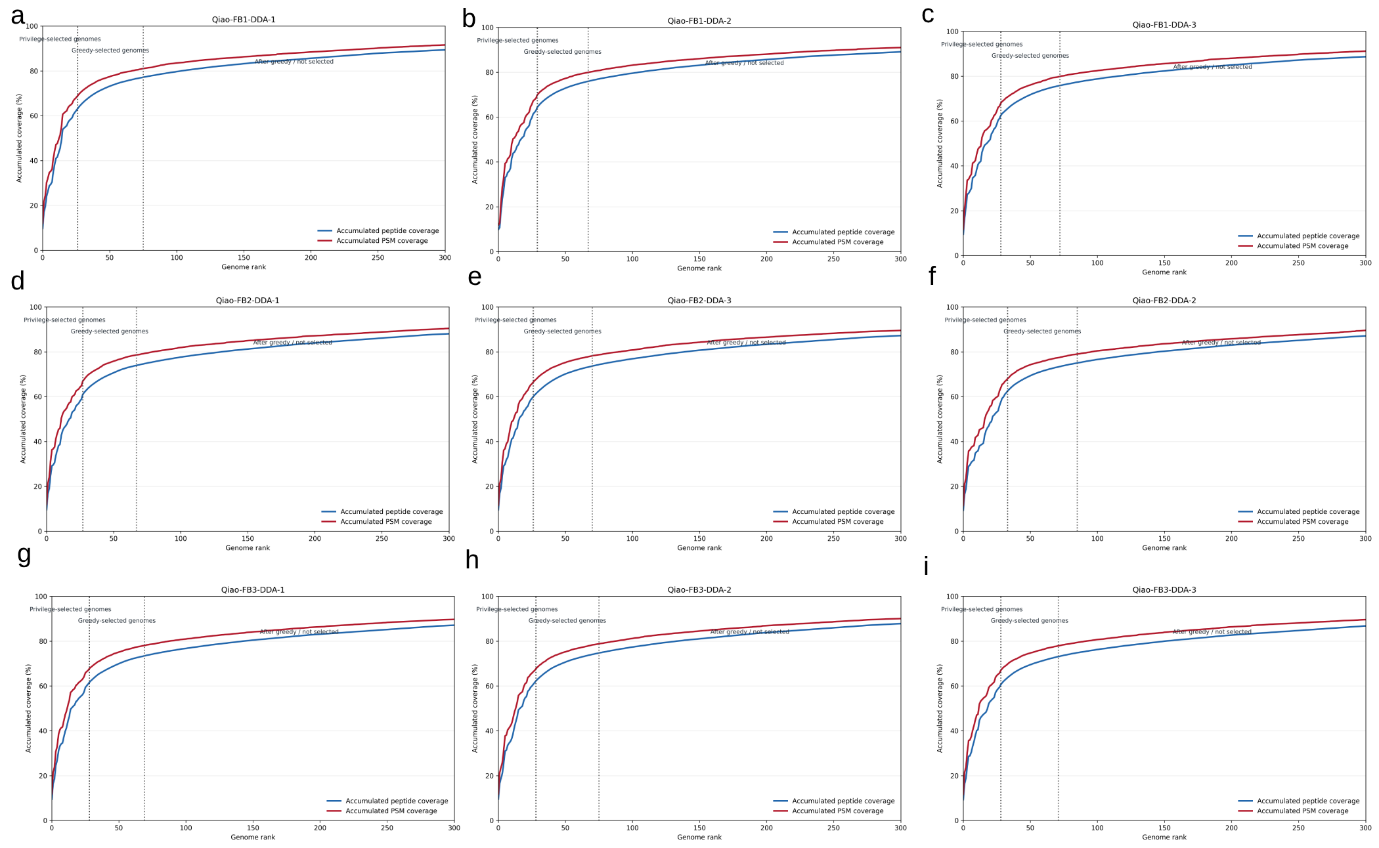

**Figure S12. Marker-peptide coverage and genome selection behaviour in Qiao_fecal_DDA samples by the Marker4 comprehensive strategy.** This dataset contains 9 raw files. Each DDA raw file is shown separately because stochastic precursor selection can lead to run-specific peptide evidence and genome-selection behaviour.

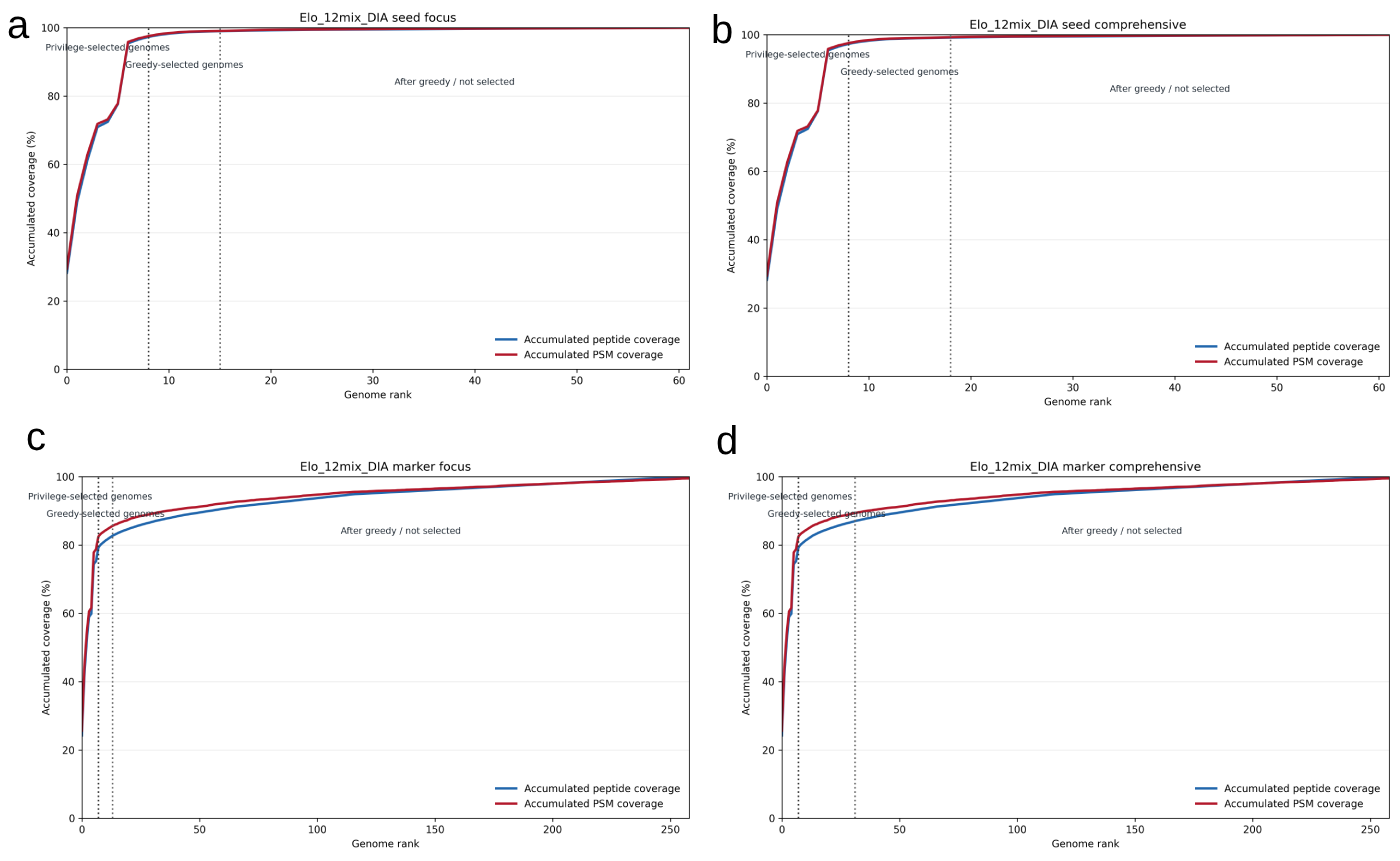

**Figure S13. Marker-peptide coverage and genome selection behaviour in Elo_12mix_DIA samples.** The four strategies are (a) seed focus; (b) seed comprehensive; (c) marker focus and (d) marker comprehensive, respectively. For DIA analyses, raw files were combined for genome selection in the pooled mode shown here; MetaPilot can also run DIA genome selection separately for each raw file.

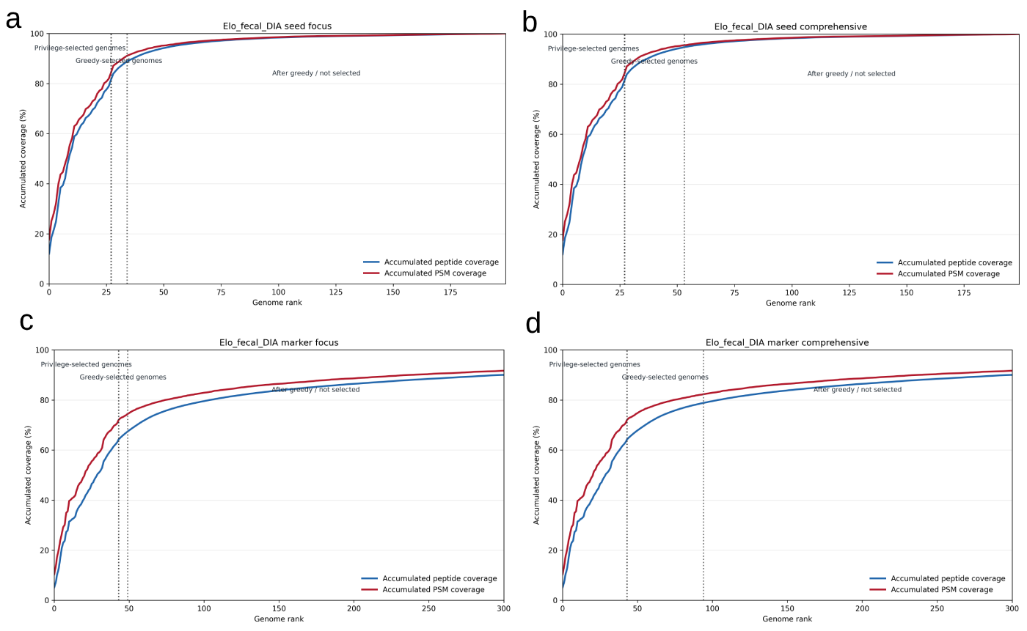

**Figure S14. Marker-peptide coverage and genome selection behaviour in Elo_fecal_DIA samples.** The four strategies are (a) seed focus; (b) seed comprehensive; (c) marker focus and (d) marker comprehensive, respectively. For DIA analyses, raw files were combined for genome selection in the pooled mode shown here; MetaPilot can also run DIA genome selection separately for each raw file.

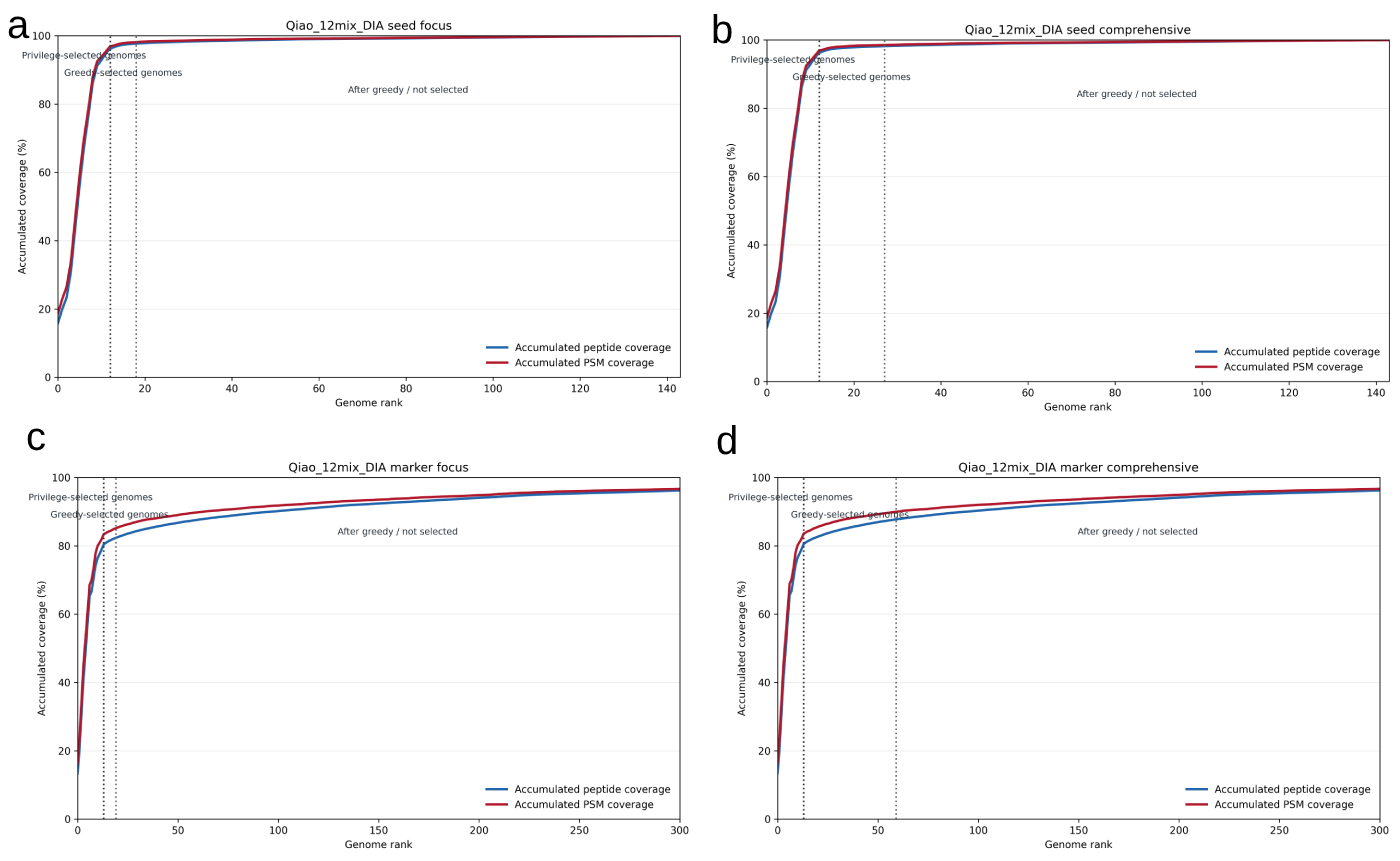

**Figure S15. Marker-peptide coverage and genome selection behaviour in Qiao_12mix_DIA samples.** The four strategies are (a) seed focus; (b) seed comprehensive; (c) marker focus and (d) marker comprehensive, respectively. For DIA analyses, raw files were combined for genome selection in the pooled mode shown here; MetaPilot can also run DIA genome selection separately for each raw file.

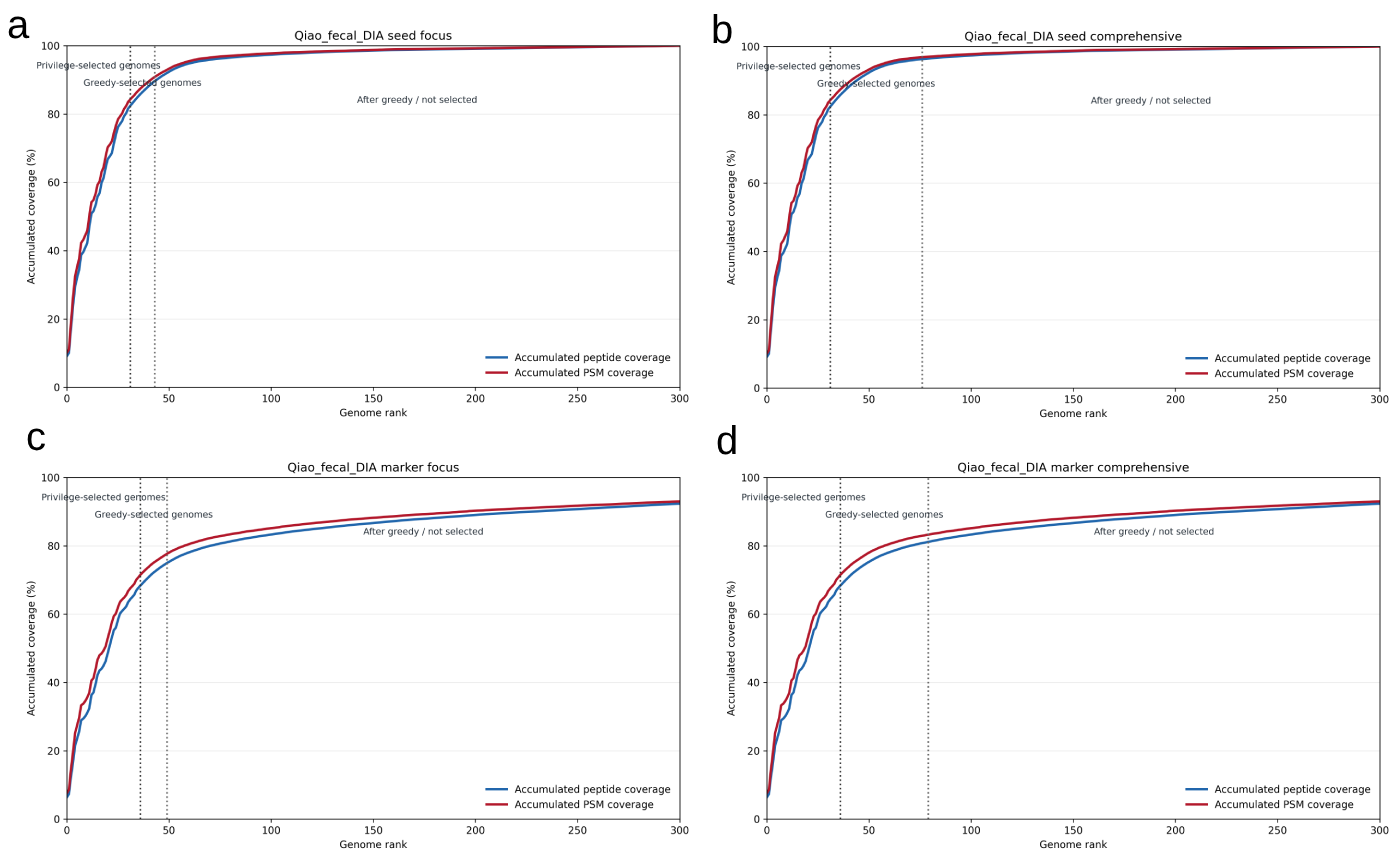

**Figure S16. Marker-peptide coverage and genome selection behaviour in Qiao_fecal_DIA samples.** The four strategies are (a) seed focus; (b) seed comprehensive; (c) marker focus and (d) marker comprehensive, respectively. For DIA analyses, raw files were combined for genome selection in the pooled mode shown here; MetaPilot can also run DIA genome selection separately for each raw file.

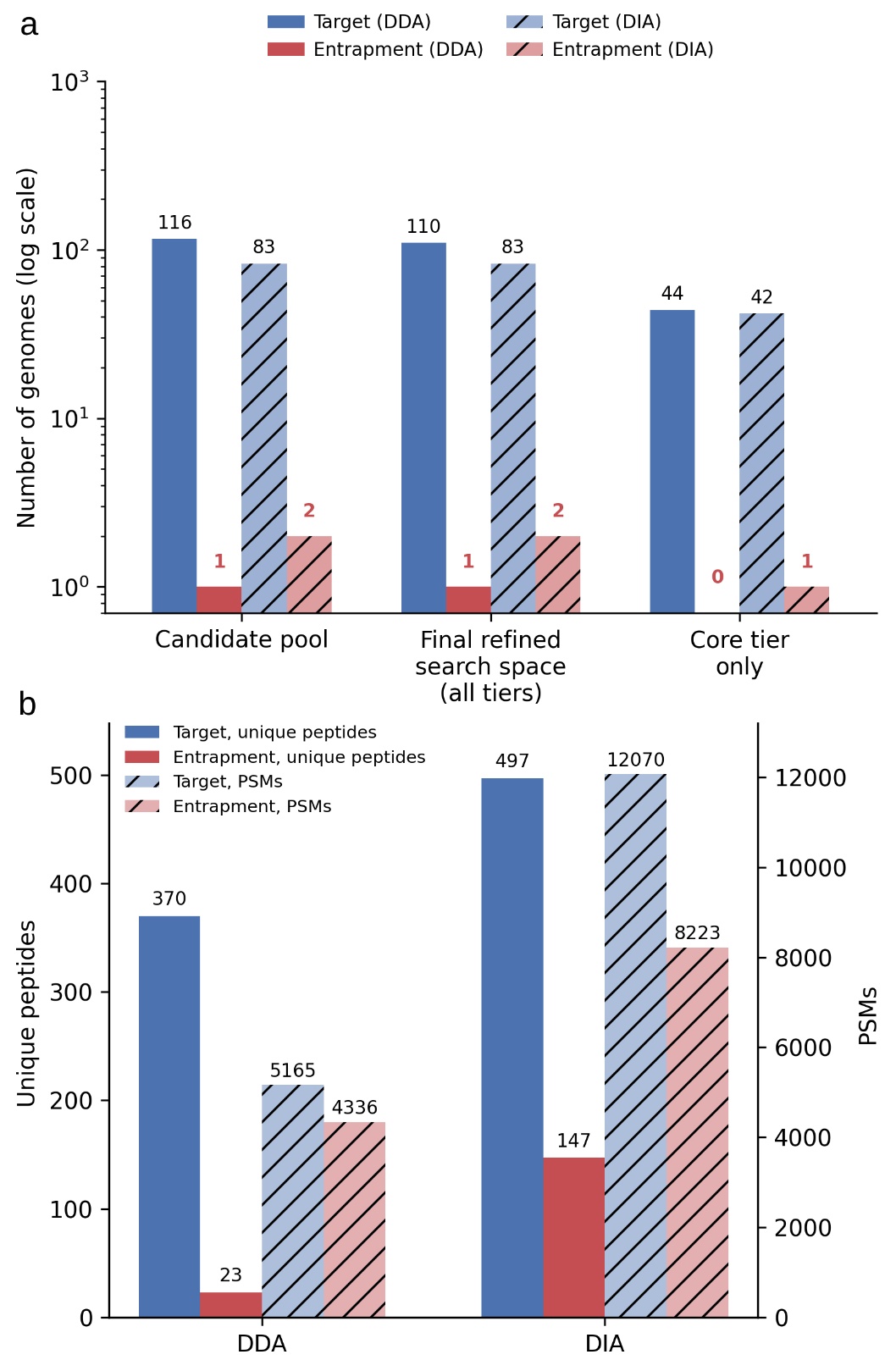

**Figure S17. (a)** Entrapment genomes are excluded from the refined search space at each stage of adaptive genome selection. Number of genomes (log scale, y-axis) classified as target (4,744-genome UHGG catalogue) or entrapment (562 tomato-rhizosphere genomes merged into the combined 5,306-genome search catalogue) at three successive stages of the MetaPilot pipeline: the genome-level significance-tested candidate pool, the final refined search space after greedy/knee-based selection (all confidence tiers combined), and the subset of genomes assigned to the highest-confidence "core" tier. Solid bars, DDA; hatched bars, DIA; blue, target genomes; red, entrapment genomes. Numeric labels give exact genome counts per bar. **(b)** The target catalogue's *Pseudomonas aeruginosa* representative independently outperforms the surviving entrapment genome *Pseudomonas paraeruginosa* in both acquisition modes. Unique peptide counts (solid bars, left axis) and PSM counts (hatched bars, right axis) for the target genome MGYG000002463 (*Pseudomonas aeruginosa*, 72 constituent genomes) and the entrapment genome MGYG000517330 (*Pseudomonas paraeruginosa*) — the only entrapment genome retained in the core confidence tier in DIA (tail tier in DDA) — shown for both acquisition modes. Blue, target; red, entrapment.

### Supplementary Table 2

| **Dataset** | **Published workflow** | **Database/library strategy** | **DDA dependency** |
| --- | --- | --- | --- |
| Qiao_faecal_DIA | directDIA-HAPs | DIA searched against HAPs-filtered database | No matched DDA library |
| Qiao_faecal_DIA | directDIA-Combined UMGS | DIA searched against combined UMGS HAPs-filtered database | No matched DDA library |
| Qiao_faecal_DIA | directDIA-MG | DIA searched against metagenomic sequencing-derived database | No matched DDA library |
| Qiao_faecal_DIA | hyblibDIA-HAPs | Hybrid library using DDA+DIA evidence with HAPs-filtered database | Yes |
| Qiao_faecal_DIA | hyblibDIA-Combined UMGS | Hybrid library using DDA+DIA evidence with combined UMGS HAPs-filtered database | Yes |
| Qiao_faecal_DIA | hyblibDIA-MG | Hybrid library using DDA+DIA evidence with metagenomic database | Yes |

### Supplementary Data

| Supplementary Data 1 | Microbiome peptide, protein, genome and function lists from the Elo_12mix samples, each DDA dataset contains 4 data tables (marker2_focus, marker2_comprehensive, marker4_focus and marker4_comprehensive); each DIA dataset contains 7 data tables (marker_focus, marker_comprehensive, marker_deep, seed_direct, seed_focus, seed_comprehensive and seed_deep) |
| --- | --- |
| Supplementary Data 2 | Microbiome peptide, protein, genome and function lists from the Elo_fecal samples, each DDA dataset contains 4 data tables (marker2_focus, marker2_comprehensive, marker4_focus and marker4_comprehensive); each DIA dataset contains 7 data tables (marker_focus, marker_comprehensive, marker_deep, seed_direct, seed_focus, seed_comprehensive and seed_deep) |
| Supplementary Data 3 | Microbiome peptide, protein, genome and function lists from the Qiao_12mix samples, each DDA dataset contains 4 data tables (marker2_focus, marker2_comprehensive, marker4_focus and marker4_comprehensive); each DIA dataset contains 7 data tables (marker_focus, marker_comprehensive, marker_deep, seed_direct, seed_focus, seed_comprehensive and seed_deep) |
| Supplementary Data 4 | Microbiome peptide, protein, genome and function lists from the Qiao_fecal samples, each DDA dataset contains 4 data tables (marker2_focus, marker2_comprehensive, marker4_focus and marker4_comprehensive); each DIA dataset contains 7 data tables (marker_focus, marker_comprehensive, marker_deep, seed_direct, seed_focus, seed_comprehensive and seed_deep) |
| Supplementary Data 5 | Microbiome peptide list from the metaExpertPro Guangzhou Nutrition and Health Study (GNHS) dataset |
| Supplementary Data 6 | Microbiome protein, genome and function lists from the metaExpertPro Guangzhou Nutrition and Health Study (GNHS) dataset |
| Supplementary Data 7 | Host peptide and protein lists from the metaExpertPro Guangzhou Nutrition and Health Study (GNHS) dataset |
| Supplementary Data 8 | Microbiome COG, NOG, KEGG KO, GO, EC and BRITE lists from the metaExpertPro Guangzhou Nutrition and Health Study (GNHS) dataset |
| Supplementary Data 9 | MGYG000002293 (*Prevotella sp900557255/ Segatella copri*) peptide and protein lists from the metaExpertPro Guangzhou Nutrition and Health Study (GNHS) dataset |
| Supplementary Data 10 | Microbiome peptide list from the colonic-content dataset |
| Supplementary Data 11 | Microbiome protein, genome and function lists from the uMetaP colonic-content dataset |
| Supplementary Data 12 | Microbiome COG, NOG, KEGG KO, GO, EC and BRITE lists from the uMetaP colonic-content dataset |
| Supplementary Data 13 | Host peptide and protein lists from the uMetaP colonic-content dataset |
